# Label-free tracking and mass measurement of single proteins on lipid bilayers

**DOI:** 10.1101/2021.04.08.438951

**Authors:** Eric D. B. Foley, Manish S. Kushwah, Gavin Young, Philipp Kukura

## Abstract

We introduce dynamic mass photometry, a method for label-free imaging, tracking and mass measurement of membrane-associated proteins. Our method enables quantitative studies of their mobility, membrane affinity and interactions at the single molecule level. Application to the membrane remodelling GTPase dynamin1 reveals heterogeneous mixtures of oligomers suggesting that the fundamental building block for oligomerisation is a dimer, challenging current tetramer-centric models. Dynamic mass photometry has the ability to transform our approach to studying biomolecular mechanisms in and on lipid bilayers.

## Main

Integral membrane proteins (IMPs) and membrane-associated proteins (MAPs) are essential for a variety of cellular processes such as signalling and vesicular trafficking, making them therapeutic targets^1,2^, with their function often relying on homo- and hetero-oligomerisation^3,4^. This complexity, combined with the need for lipid bilayers, makes it particularly challenging to accurately characterise the stoichiometries and kinetics of the biomolecular interactions underlying IMP and MAP function and regulation. Advances in single molecule fluorescence-based microscopy methods^5,6^ have enabled *in vivo* and *in vitro* investigations into the interactions of IMPs, such as dimerisation of GPCRs^7,8^, nano-clustering^9^, and of MAPs, such as the coordination of Min proteins in bacterial cell division^10^, and the mechanism of amyloid-ß plaque formation on cell membranes, which is associated with Alzheimer’s disease^11^. The main challenges in fluorescence-based methods, however, arise from quantitative uncertainties caused by incomplete labelling of the sample, photochemical and photophysical effects such as photoblinking, photobleaching and quenching, and the distinct labelling required to detect multiple species simultaneously. These limitations have made it challenging to accurately quantify processes such as membrane (un)binding of MAPs and the dynamics and stoichiometries of protein-protein interactions for both MAPs and IMPs. While numerous approaches aimed at molecular subunit counting exist^12–14^, the analysis and interpretation of the resulting oligomeric distributions is complicated and the number of heterogeneous species that can be detected simultaneously remains limited. Given the critical functional importance of homo- and hetero-oligomeric interactions for membrane-associated processes, there is an urgent need for a quantitative and dynamic approach capable of complementing the information accessible from existing methods.

Mass photometry (MP) is a label-free method based on interferometric scattering microscopy that detects single biomolecules in solution and measures their mass with an overall mass accuracy and resolution of 1-2% and 20 kDa, respectively^15^. These capabilities enable the quantification of protein-protein interactions with sufficient sensitivity to accurately determine stoichiometries and rates of reactions^16^. As such, MP could be ideally suited to address the shortcomings of existing fluorescence-based techniques for *in vitro* applications of IMPs and MAPs. Existing implementations of MP, however, rely on stationary binding of individual molecules to a surface, usually a microscope coverslip. By averaging images taken before a binding event and subtracting them from averaged images taken after a binding event, the signal due to surface roughness is removed and the shot noise is lowered sufficiently to detect individual molecules binding to the surface^17–19^. Here, we show that MP can be extended to *in vitro* studies of individual protein complexes on supported lipid bilayers (SLBs).

To explore the capabilities of MP on an SLB (60/40 PC/PS, see methods), we chose the 100 kDa MAP dynamin1-WT (WT). Its function relies on (dis)assembly on lipid bilayers (Fig. 1a), with our current understanding of the underlying molecular mechanisms of dynamin relying entirely on assumptions based on structural information and bulk behaviour. As expected, raw MP images exhibited background from the roughness of the microscope coverslip. By implementing a sliding median background subtraction, we could obtain nearly shot noise-limited background and diffraction-limited features arising from individual WT complexes diffusing on the SLB (Supplementary Fig. 1, Supplementary Movie 1). The sliding median correction involves estimating the background from the temporal median of a series of frames around each frame of interest (see methods). Importantly, this approach avoids the convolution of scattering contrast and particle motion inherent to the background subtraction used in standard MP, and reduces the imaging background at equivalent imaging speeds due to the larger number of frames contributing to the background image (Supplementary Fig. 1–2).

**Figure 1.**
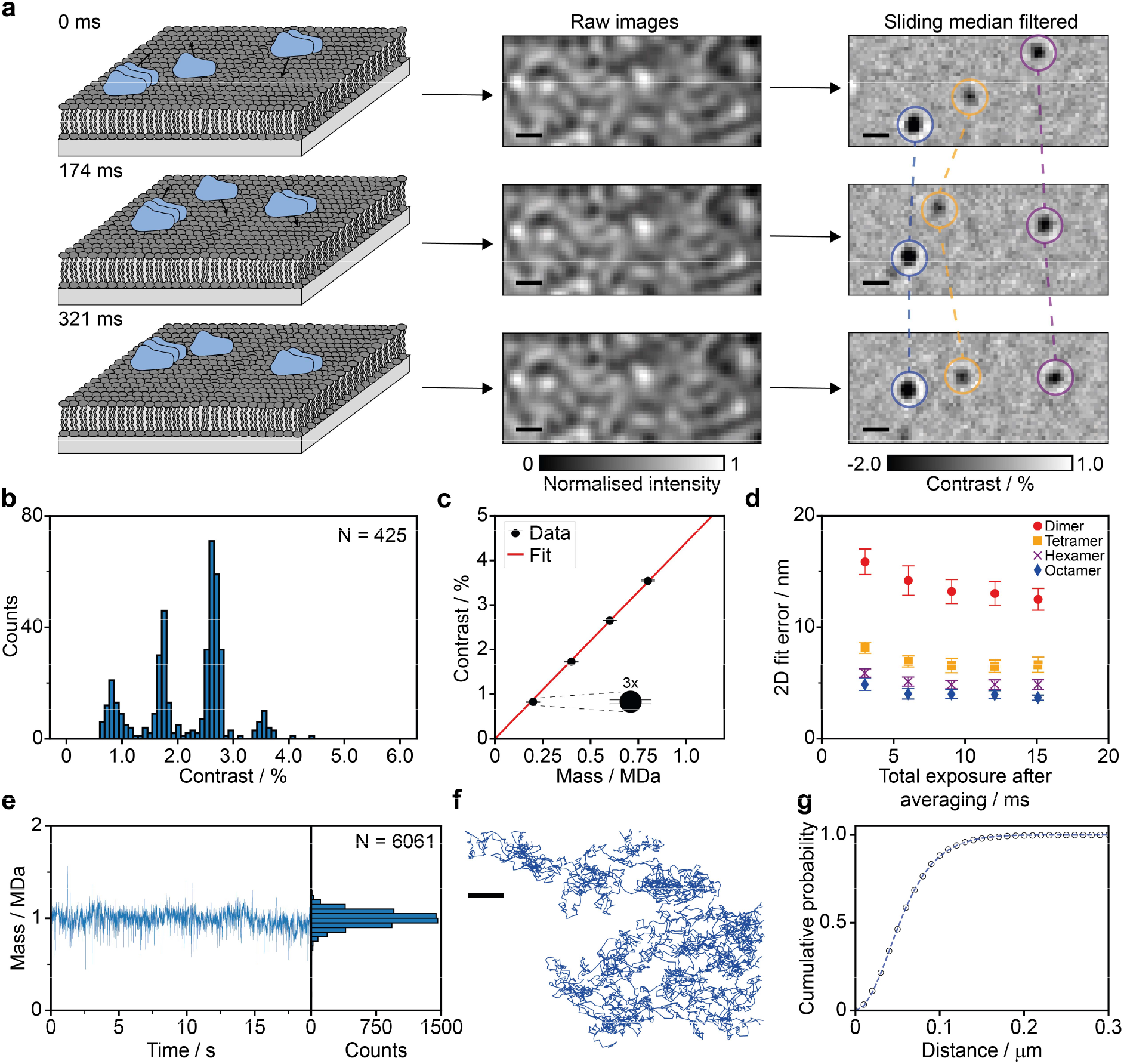
Principle and performance of dynamic mass photometry. **(a)** Schematic of a dynamic mass photometry measurement with protein complexes diffusing on a supported lipid bilayre. Images were acquired at 331 Hz and then processed with a sliding median filter (see methods), which revealed individual protein complexes on the bilayer as diffraction-limited spots. **(b)** Histogram of mean contrasts of each particle trajectory after length filtering detected in a dynamic MP movie (4 min) of WT diffusing on an SLB. **(c)** Contrast vs mass calibration curve of the contrast distribution in **b** yielding a contrast to mass ratio of 4.40 %/MDa. Error bars are plotted along the contrast axis and correspond to the standard error of the mean of the Gaussians that were fitted to the contrast histogram. **(d)** 2D localisation error of our PSF fitting procedure of dynamin dimers (red circles), tetramers (orange squares), hexamers (purple crosses) and octamers (blue diamonds) plotted as a function of effective exposure time. Errors are reported as the combined standard deviation of the error in x and y. **(e)** Mass trace and histogram of a 19 s WT-dynamin decamer trajectory. **(f)** Corresponding particle trajectory. **(g)** Corresponding cumulative probability of particle displacements during 1 frame (t = 3 ms) and fits to a two-component model (Eq. 4). Scale bars: 500 nm.

For the chosen system, particles exhibited clearly differing signal intensities (Fig. 1a, Supplementary Movie 1). Filtering for trajectories that remained bound to the membrane for at least 50 frames, corresponding to a residence time of 151 ms (Supplementary Fig. 3), and plotting the mean contrast of the remaining 600 trajectories revealed a contrast distribution with equally-spaced peaks, as expected for different oligomeric states of the same protein (Fig. 1b). The contrast values of these particles increased linearly with mass (Fig. 1c) and matched well with the contrasts measured for dimers, tetramers, hexamers and octamers in standard MP of the 90 kDa dynamin mutant, ΔPRD (Supplementary Fig. 4), demonstrating that dynamic MP can simultaneously image, track and measure the mass of diffusing biomolecular complexes. Additionally, the oligomeric distribution of WT on the SLB was strikingly different from that measured in solution by standard MP (Supplementary Fig. 5), with many higher oligomeric species present on the SLB.

Localisation precision and imaging speed are key performance parameters for single particle tracking, and determine how much information can be extracted from individual trajectories. The nature of the background subtraction used in dynamic MP prevented us from assessing the localisation precision by repeatedly measuring the location of surface-immobilised particles, as is commonly done in fluorescence-based methods. Nevertheless, we could estimate the localisation precision by extracting the error of our point spread function (PSF) fitting procedure (Fig. 1d). At our imaging speed of 331 Hz (3 ms total exposure), the fit error for WT dimers (200 kDa particles) was 16 nm, which compares well with the localisation precision in single molecule fluorescence imaging at similar speeds (18 nm)^20^, and improved with increasing mass (8, 6 and 5 nm for WT tetramers, hexamers and octamers, respectively). Localisation errors improved by up to 20% when lowering the effective imaging speed to 110 Hz (see methods), beyond which there was no further improvement. On average, we found that the slope of the contrast vs mass calibration in dynamic MP was 10-15% lower than in standard MP, where particles remain stationary. This trend became more pronounced as we lowered the effective imaging speed from 331 Hz to 66 Hz, resulting in a drop in contrast precision of 20% and a further 15% decrease in particle contrast (Supplementary Fig. 6). As such, we attribute these effects to motion blurring of the PSFs, which is caused by particle movement during image acquisition, thus resulting in decreased particle contrast and diminished improvements in localisation precision at slower imaging speeds. We thus chose to image at 331 Hz to minimise the effects of motion blurring. As a result, however, we were unable to detect WT monomer particles on the SLB, and in some cases it was difficult to distinguish dimer particles from background noise.

As dynamic MP is not subject to photobleaching, the time limit on observing particle trajectories is in principle only determined by how long particles remain bound to the membrane or within the field of view. As a result, we could obtain trajectories consisting of up to 6000 frames (20 s) with continuous observation, consistent localisation precision and mass measurement (Fig. 1e-f and Supplementary Movie 2). From this data, we could compute the diffusion coefficient by fitting multiple-mobility models to the cumulative probability distribution of particle displacement (Supplementary Fig. 7a) during a defined lag time, *t* (Eq. 3–5)^5^. For the WT decamer particle in Fig. 1e-f, a two-component fit was determined to be most suitable (see methods), and revealed major and minor diffusion coefficients of D_1_ = 0.58 μm^2^ s^−1^ and D_2_ = 0.22 μm^2^ s^−1^ with relative weightings of 0.56 and 0.44, respectively, using t = 3 ms (Fig. 1g).

By applying this approach to dynamic MP measurements of 20 nM WT, we were able to measure the diffusion coefficients of different oligomeric species, which were resolved in the mass distribution with a full width at half maximum of <50 kDa (Fig. 2a). Using the smallest possible lag time (t = 3 ms), over 95% of species only exhibited one type of diffusive behaviour (Supplementary Fig. 7b), as expected for simple Brownian motion. Further repeat measurements with WT (Supplementary Fig. 8-10) and ΔPRD (Supplementary Fig. 11-12), which is more oligomerisation-prone than WT^21^, revealed a reproducible inverse proportionality of the diffusion coefficients with the number of oligomeric subunits (Fig 2b). Given that the diffusion of membrane-bound proteins has been reported to primarily depend on their contact area with the SLB and the number of bound lipids^22–24^, our results suggest that the contact between the SLB and the oligomers of WT and ΔPRD in the range observed here increases linearly with oligomeric state. Additionally, we observed an increase in calculated diffusion coefficients of all oligomeric species when increasing the lag time from 3 to 12 ms (Supplementary Fig. 13a), most likely caused by the dynamic error in our dynamic MP measurements due to a combination of the relatively fast particle motion combined with nanometre localisation precision^25^. At longer lag times there was little change in the diffusion coefficients, again confirming that dynamin undergoes Brownian motion on the SLB on the time scales observed in this study.

**Figure 2.**
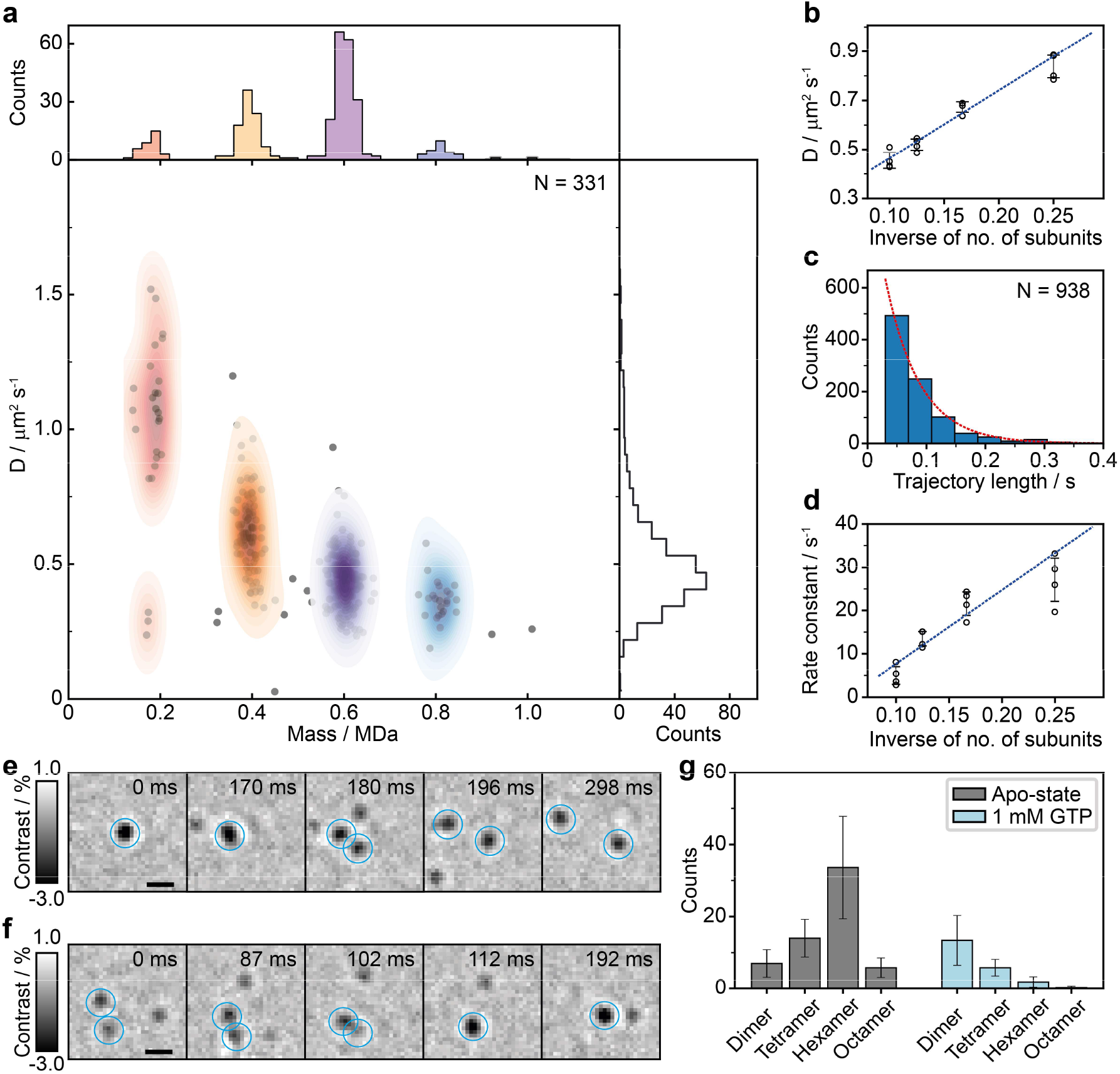
Oligomeric properties and dynamics of dynamin diffusing on a supported lipid bilayer. **(a)** Major diffusion components vs mean trajectory mass for a 4 min movie of WT-dynamin (20 nM solution concentration). **(b)** Major diffusion components of each oligomeric species vs the inverse of the number of oligomeric subunits from 4 replicate dynamic MP measurements of ΔPRD (10-20 nM, 4-5 min each) and a corresponding weighted linear fit. **(c)** Histogram of SLB residence times of ΔPRD hexamer particles from one of the dynamic MP movies with a fit to a 1-component exponential distribution (appropriately scaled here for display) yielding a dissociation rate constant of 17.3 s^−1^. **(d)** Dissociation rate constants of each oligomeric species vs the inverse of the number of oligomeric subunits from the same experiments used in **b** and a corresponding weighted linear fit. **(e)-(f)** Examples of a dissociation and an association event. **(g)** Effect of GTP addition on the oligomeric distribution of 10 nM WT (5 replicates of 1 min dynamic MP movies before and after GTP addition). Error bars in all figures correspond to the standard deviation determined from the indicated number of replicate measurements. Scale bar: 500 nm.

We were also able to quantify SLB residence times and their dependence on the oligomeric state. We found that the distributions of residence times of both WT and ΔPRD oligomers were well described by an exponential model (Fig. 2c), as expected for a first-order process, from which we could extract the dissociation rate for each oligomeric species (Supplementary Fig. 14-15). Similar to the diffusive behaviour, the dissociation constants were inversely proportional to the number of subunits (Fig. 2d, Supplementary Fig. 16), further confirming that SLB contact increases linearly with oligomeric state. Additionally, we found instances of dynamin oligomers dissociating into subunits (Fig. 2e Supplementary Movie 3) and vice versa (Fig. 2f, Supplementary Movie 4) on the SLB. Such events were rarely observed in our systems (in less than 0.1% of trajectories), suggesting that they tend to happen on timescales longer than the SLB residence times for dynamin in its apo-state (<200 ms on average). Upon addition of GTP, which is essential to dynamin function and disassembly from the membrane, we found that the overall particle density of WT, in particular the proportion of large oligomers, immediately decreased (Fig. 2g and Supplementary Fig. 17), in line with GTP hydrolysis-induced disassembly^26^ and unbinding of dynamin polymers.

These results demonstrate accurate and resolved mass measurement of proteins on lipid bilayers at the single molecule level, together with quantification of key parameters such as oligomeric distributions, residence times and diffusion coefficients, and observation of (dis)association events. Our observation of dynamin dimers, tetramers, hexamers, octamers and decamers strongly suggests that dynamin oligomerisation proceeds by dimer addition, rather than being based in tetrameric subunits^27^, which could have implications for our understanding of disease-related mutations. Use of cushioned or suspended lipid bilayers^28^ will expand the demonstrated capabilities to integral membrane proteins, and interactions between them, and with soluble proteins, transforming our capabilities in assessing the biophysical properties of this important class of biomolecules.

## Methods

### Stocks, reagents, and instruments

HEPES (H3375), KCl(P9541), Imidazole (I5513), DL-Dithiothreitol (DTT: D9779), Magnesium Chloride hexahydrate (M2670), Glycerol (G5516), Isopropyl-β-D-thiogalactoside (IPTG: I6758), Ampicillin (59349), Chloramphenicol (C0378), Protease inhibitor cocktail (4693159001; Roche), HiTrap^®^TALON^®^ cobalt columns (GE28-9537-67), StrepTrap™ columns (GE28-9075-48), Grace Bio-Labs reusable CultureWell™ silicon gaskets (GBL103280), and Desthiobiotin (71610-3) were purchased from Merck Life Science UK Limited. 1,2-dioleoyl-sn-glycero-3-phosphocholine (DOPC; 850375P) and 1,2-dioleoyl-sn-glycero-3-phospho-L-serine (sodium salt) (DOPS; 840035P) were purchased from Avanti Polar Lipids. GTP (NU-1012) and GppNHp - Tetralithium salt (GMPPNP) (NU-401) were purchased from Jena Biosciences. BL21-CodonPlus and Terrific Broth were purchased from Agilent Technologies and Fisher Scientific, respectively. The plasma cleaner, and ultrasonicator are from Diener electronic and Sonics & Materials, Inc, respectively. The Dynamin1 wild type (WT) bacterial expression vector was generously provided by Prof. Thomas Pucadyil (IISER Pune, India). ΔPRD (Dyn1_1-746_) was generated in the lab using standard cloning procedures and confirmed using DNA sequencing.

HEPES, KCl, MgCl_2_, and Imidazole, stocks were prepared in Milli-Q^®^ water (18.2 MΩ·cm) (Milli-Q) and filtered using a 0.2 μm filter. Ampicillin, IPTG and DTT were prepared in degassed Milli-Q, aliquoted and stored at −20°C. Chloramphenicol stock were prepared in ethanol, aliquoted and stored in −20. GMPPNP (50 mM) was dissolved in degassed 20 mM HEPES (pH = 7.4). GTP solution and GMPPNP stocks were aliquoted and stored at −20°C. DOPC (25 mM) and DOPS (10 mM) stocks were prepared using chloroform and aliquoted and stored at −20°C under nitrogen.

### Protein expression and purification

Bacterial expression vectors for WT and ΔPRD were transformed in BL21-CodonPlus and grown until OD_600_ = 0.6 in Terrific Broth containing ampicillin (100 μg ml-1) and chloramphenicol (35 μg ml^−1^) at 37°C. Protein production was induced with 0.1 mM IPTG, and bacterial cultures were grown for 12 hours at 18°C. Cells were harvested, and bacterial pellets were stored at −80°C. For purification, bacterial pellets were thawed on ice and dissolved in lysis buffer (20 mM HEPES, pH = 7.4, 300 mM KCl and 20 mM Imidazole) containing one tablet of protease inhibitor cocktail. Cells were lysed using a cell homogenizer, and lysate was spun at 20,000 g for 20 min at 4°C to remove insoluble cell debris. The supernatant was applied to a 5 ml HiTrap^®^TALON^®^ column. The column was subsequently washed with 60 ml lysis buffer, followed by 60 ml wash buffer (20 mM HEPES, pH = 7.4, 150 mM KCl and 20 mM Imidazole) to remove non-specifically bound protein. The protein was eluted with elution buffer (20 mM HEPES, pH = 8, 150 mM KCl, 200 mM Imidazole) and eluted protein was applied to a 5 ml StrepTrap™ column, which was pre-equilibrated with elution buffer, at 2 ml min^−1^. The column was washed sequentially with 50 ml strep-1 buffer (20 mM HEPES, pH = 7.4, 300 mM KCl), 50 ml strep-2 buffer (20 mM HEPES, pH = 7.4, 300 mM KCl, 1 mM DTT), 50 ml strep-3 buffer (20 mM HEPES, pH = 7.4, 150 mM KCl, 1 mM DTT). The protein was eluted with strep-elution buffer (20 mM HEPES, pH = 7.4, 150 mM KCl + 1mm DTT+ 10% glycerol+ 2.5 mM Desthiobiotin), flash frozen and stored in −80°C until use.

To prepare the protein for dynamic MP experiments, WT or ΔPRD were dialysed overnight at 4°C in degassed and pre-chilled HKS-150 buffer (20 mM HEPES, pH = 7.4, 150 mM KCl) containing 1 mM DTT. After dialysis, the protein was spun at 20,000 g for 30 minutes at 4°C to remove aggregates. The supernatant was collected, and the concentration was estimated by measuring absorbance at OD_280_ nm and using molar extinction coefficients of 62,800 M^−1^ cm^−1^ and 54,300 M^−1^ cm^−1^ for WT and ΔPRD, respectively. Proteins were stored on ice during use.

### Supported lipid bilayer preparation

To prepare SLBs, we used sonicated liposomes consisting of a 60/40 molar ratio of DOPC/DOPS. Liposomes were prepared by adding DOPC and DOPS stock solutions into a clean glass tube at the required molar ratio (500 μM total lipid concentration) followed by drying via rotary evaporation under a constant nitrogen stream, followed by further drying under vacuum for 1 hour at room temperature to remove remnants of organic solvent. 500 μl of freshly prepared HKS-150 (for details, see buffer stock preparation for bilayers) was added to the dried lipids and the mixture was covered with parafilm (Fisher Sci.) and incubated at 50°C in a water bath for 1 hour with intermittent mixing and stored overnight at room temperature. Before sonication, the resulting liposome mix was transferred into a 1.5 ml Eppendorf tube and kept in ice-water. For sonication, a 3 mm probe was used at 25% amplitude with a 1 s on, 3 s off sonication cycle for 10 minutes. Sonicated liposomes were then spun at 20,000 g for 30 minutes to remove debris and other insoluble contents. Supernatant was collected in a new Eppendorf tube, stored in the fridge, and used within 3 days of sonication.

To prepare fluid SLBs, glass coverslips were cleaned by sonication in Milli-Q water for 5 min, then isopropanol (5 min) and again water (5 min) in an ultrasonic bath and then dried using a nitrogen stream and stored in a dry place until use. Before SLB preparation, coverslips were treated with oxygen plasma for 8 minutes using a plasma cleaner and used immediately. Long-term exposure to air (>1-2 h) or plasma cleaning for only 2-3 minutes led to either incomplete surface coverage of the SLBs or less fluid SLBs (characterised by significantly more than 5% of molecules displaying a second diffusion component in dynamic MP experiments).

For SLB preparation, silicon gaskets were rinsed with Milli-Q water, then isopropanol, and again Milli-Q water, and dried under a nitrogen stream. Cleaned gaskets were placed on the freshly plasma cleaned coverslips and 30 μl of freshly reconstituted HKS-150 containing 1.7 mM MgCl_2_ buffer was added to the gasket. 20 μl of sonicated liposomes were then added to the gasket, followed by mixing with a micropipette and incubation at room temperature for at least 30 minutes, and left undisturbed at room temperature until further use. Inclusion of 1 mM MgCl_2_ at the step of bilayer formation proved crucial to form fluid and continuous SLBs. Before use of the SLBs, 15 μl of fresh HKS-M (20 mM HEPES, pH = 7.4, 150 mM KCl, 1 mM MgCl_2_) was added to the gasket to account for the loss of volume due to evaporation. Subsequently, 40 μl of solution in the gasket was replaced with 40 μl of fresh HKS-M to wash away excess liposomes and micro-particles. This process was repeated at least 10 times and gaskets were inspected under the mass photometer for the presence of impurities and micro-particles further washes were conducted if necessary. After washing, HKS-M was replaced with reaction buffer (20 mM HEPES, pH = 7.4, 100 mM KCl, 1 mM MgCl_2_) by further serial dilution by replacing 40 μl gasket solution with reaction buffer (6 times) and inspected for presence of any floating particles in the MP setup.

### Buffer preparation

Due to its label-free nature, dynamic MP is sensitive to the presence of scattering impurities. These impurities can integrate into the SLBs and introduce defects or remain stuck to the SLBs in the field of view. To prevent this, buffer stocks were prepared extremely carefully. HEPES (1 M, pH = 7.7), KCl (3 M) and MgCl_2_ (1 M) stocks were prepared in degassed Milli-Q water and filtered twice with 0.2 μm filter and stored at 4°C until use. Buffers were reconstituted on the day of the dynamic MP experiments and prepared in degassed Milli-Q water. The required amounts of stocks were aliquoted in Milli-Q and mixed by vortex. Buffers were replaced every 2-3 hours to decrease the influence of reactive oxygen species during dynamic MP measurements.

### Mass photometry setup

All data except for the trajectories in Fig. 1a, 1e-g and 2e-f were acquired on a Refeyn OneMP mass photometer with a 10.8 × 2.9 μm^2^ (128 × 35 pixels) field of view. The microscope used to acquire the data in Fig. 1a, 1e-g and 2e-f was custom-built with a 9.4 × 6.2 μm^2^ field of view and is similar to that described in previous work from our group^18^. The custom-built setup is illustrated in Supplementary Fig. 18 and the differences to the setup used in previous work are highlighted in the figure caption.

### Data acquisition

To acquire dynamic MP movies, the prepared coverslips with SLBs (contained by a silicon gasket filled with buffer) were placed on the sample stage to optimise the focus of the microscope. After locking the focus of the microscope, WT or ΔPRD was added to the SLB by replacing 3-6 μl of buffer from silicon gasket with 3-6 μl of 200 nM WT or ΔPRD (in HKS-100) solution and mixing well with a micropipette to achieve a final concentration of 10-20 nM. Data acquisition was then started within 10 seconds of protein addition. Images were collected at 994 Hz over a period of 5-10 minutes and saved after binning blocks of 3×3 pixels and binning frames into groups of 3, resulting in an effective frame rate of 331 Hz and final pixel size of 70.3 nm. This level of frame averaging was chosen because it was the fastest effective frame rate that allowed for the detection of WT dimer particles (0.2 MDa), while minimising the effects of motion blurring. At lower effective frame rates (increased frame averaging), we observed a drop in particle contrast and contrast precision (Supplementary Fig. 6), which we attribute to motion blurring during frame acquisition and averaging. We note that the contrast of WT dimer partially overlaps with background noise, which likely caused the contrast to plateau with decreasing frame rate instead of further decreasing as for the other oligomeric species. Due to this overlap, we did not include WT and ΔPRD dimer in Fig. 2b, d and Supplementary Fig. 10, 16.

Data acquisition was briefly paused (~5-10 s) once every minute (20,000 frames) to readjust the microscope focus to account for drift over time before resuming acquisition. This was repeated several times to obtain sets of 4-8 movies (1 min each) for that were then combined into one data set for each SLB and sample. For experiments on the effect of GTP/GMPPNP, 3-4 movies of WT (10-20 nM) were recorded as described above, after which image acquisition was briefly paused and 1.2 μl of GTP or GMPPNP (50 mM) was added (total gasket volume = 60 μl), followed by mixing to obtain a final concentration of ~1 mM before resuming acquisition to collect another 3-4 movies. The number of replicate measurements indicated in figure captions corresponds to the number of sets of movies that were taken for each protein. We used the same purified batch of WT and ΔPRD for all data collected on the OneMP setup.

### Data processing

#### Background subtraction

Dynamic MP movies were processed by treating each frame with a sliding median background subtraction algorithm. In brief, each frame was divided by its ‘local median’, *i.e*. the median of a pre-defined frame interval (here 201 frames or 607 ms) centred around the frame of interest, to calculate the background-subtracted frames, F:

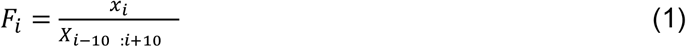

where *x*_*i*_ is the current raw frame and *X*_*i-100:i+100*_ represents the median pixel values of raw frames, from i-100 up to (and including) i+100. Each background-subtracted frame was then additionally treated with a 2D-median noise filter to remove any large dynamic background sources (*e.g.* fluctuations in illumination, if present). The window size of 201 frames for the sliding median algorithm was chosen because it was the smallest window size that did not detrimentally affect particle contrast or contrast precision (Supplementary Fig. 19). For smaller window sizes, particle contrast values and contrast precision decreased significantly, especially for larger particles that were less mobile, while larger window sizes increased processing times without an additional increase in sensitivity or performance. We anticipate that for slower moving particles (D < 0.3 μm^2^ s^−1^), it may be necessary to further increase the window size to avoid detrimental effects on performance.

#### Particle detection

Particle candidates were identified by treating each processed frame with a Laplacian of Gaussian filter that matched the size of the PSFs in our MP setups (Supplementary Fig. 20). From this filtered image two binary maps were constructed by a) applying a manually set threshold and b) applying a local maximum filter. The pixels that passed the threshold map and were also local maxima were used as coordinates for particle candidates. For each pair of candidate coordinates, a 13×13 pixel region of interest was constructed with the candidate pixel at the centre and this region of interest was passed through our PSF fitting procedure to quantify particle contrast and location. If a particle candidate was too close to an edge of the field of view to construct a 13×13 region of interest, *i.e*. within 6 pixels of an edge, it was discarded. In some cases, background noise features were identified as particle candidates and this could lead to the PSF fit converging onto a nearby particle in the region of interest, which resulted in duplicate fits. To avoid problems with trajectory linking, only the first instance of a fitted particle was kept and any duplicates were deleted.

#### Particle quantification and point spread function model

Particle candidates were quantified through least-squares minimisation of the residual between the 13×13 region of interest and our PSF model. Due to the interferometric nature of dynamic MP, we based our PSF model on the shape of a jinc function^29^ rather than its square, which is more commonly used in fluorescence-based techniques:

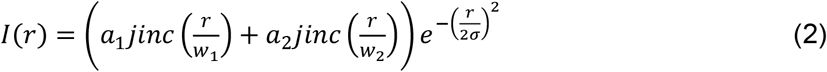

The first jinc function models the light scattered by a small particle, which is clipped by the circular objective aperture, where r is the distance from the PSF centre, *w* the width of the jinc function and a its amplitude. In MP setups a partial reflector positioned in the back focal plane helps to increase particle contrast by attenuating the light reflected by the coverslip^18^, which we account for by including a second jinc function. This combination of two jinc functions is then multiplied by a Gaussian with standard deviation σ, which is an empirical adjustment to reflect the appearance of the PSFs in our setups, which appear to have weaker outer lobes than we can account for with just jinc functions. We calibrated this PSF model using standard MP landing assay with 8-fold frame averaging compared to the dynamic MP measurements to achieve a higher signal-to-noise ratio. These landing assays were carried out within 2 hours of the dynamic MP experiments. We then extracted and saved the ratio of the amplitudes of the two jinc functions (*a*_*1*_/*a*_*2*_), the width of the first jinc function (*w*_*1*_) and the standard deviation of the Gaussian (σ). The width of the second jinc function (*w*_*2*_) is calculated through prior knowledge of the dimensions of the back aperture and partial reflector (here *w*_*2*_ = 2.27*w*_*1*_). The analysis of these landing assays was carried out in DiscoverMP software (Refeyn Ltd) and the extracted parameters used for each measurement are supplied with the raw data.

#### Trajectory linking

The successfully fitted particles were linked into trajectories using the open source python package trackpy^30^. More specifically, we used the trackpy.link_df function with a maximum search distance of 4 pixels from frame to frame and a ‘memory’ of 3 frames. The memory parameter refers to the maximum number of frames during which a feature can vanish (as a result of unsuccessful PSF fitting, for example) and reappear and still be considered the same particle. Due to this memory parameter, our linked trajectories can contain gaps of up to 3 frames in length each. To obtain accurate trajectory lengths, the missing frames were treated as trajectory points where the contrast and position could not be determined.

### Trajectory analysis

#### Trajectory filtering

Unless otherwise stated, linked trajectories were processed as described here. Only trajectories that lasted at least 151 ms (50 frames) were used for analysis, as this effectively reduced the amount of background noise features and incorrect linking, and improved contrast resolution (Supplementary Fig. 3). Additionally, particle trajectories that had coordinates within 5 pixels of the edge of the field of view were discarded to avoid artificial trajectory shortening caused by particles leaving (and sometimes re-entering) the field of view. Next, we constructed a contrast histogram for each trajectory and applied a Gaussian fit to extract the mean and standard deviation of the contrast of each trajectory. These mean trajectory contrasts were then filtered by their standard deviation to further eliminate poorly linked or noisy trajectories (Supplementary Fig. 21). For this filtering step, we used a contrast vs standard deviation trend obtained from an MP landing assay of ΔPRD on the same instrument with 8-fold frame averaging (Supplementary Fig. 21a) and applied it with an appropriate contrast offset to the trajectories obtained after length filtering (Supplementary Fig. 21b). This offset was identified by inspection to account for the additional variation caused by operating at a faster frame rate (offset = 0.0015 at 331 Hz). Examples of trajectories that were kept and rejected based on this filtering step are shown in Supplementary Fig. 21d-e. After these two filtering steps, the mean trajectory contrasts were plotted in histograms (70 bins) for WT and ΔPRD. We then used Gaussian fitting to the resulting contrast distribution to self-calibrate the data, and convert contrast to mass and allocated particle trajectories into different oligomeric states (i.e. a trajectory was identified as belonging to a particular oligomer if its mean mass was within two standard deviations of the mean mass of one of the oligomeric species; see Supplementary Fig. 8 and 11 for examples of this selection range).

#### PSF fitting error of particle locations

For every successful PSF-fitting operation, the error of the fit in x and y was estimated from the covariance of the parameters of the fit. To determine the mean fitting errors in 2 dimensions for each oligomeric species of WT, we examined the data shown in Fig. 1b-c at different effective imaging speeds, which was achieved by averaging together increasing numbers of consecutive frames (2, 3, 4 and 5, resulting in effective frame rates of 166 Hz, 100 Hz, 83 Hz and 66 Hz) of the dynamic MP movie after treatment with the sliding median filter. Trajectories were linked and filtered as described above (with incrementally increasing offsets from 0.0015-0.0023 for frame rates from 331 Hz to 66 Hz). The median PSF fitting error in x and y was then determined for each trajectory that passed the filtering steps and a Gaussian was fit to the resulting distribution of median x and y errors of the trajectories making up each oligomeric peak to extract the mean and standard deviation. The fitting errors in 2D particle positions displayed in Fig. 1d were calculated from the means of these Gaussian fits. We note that this provides a lower bound on the localisation precision of dynamic MP at our operating frame rate. Trajectory linking parameters were adjusted for decreased effective frame rates (max search distance of 4, 6, 6, 7 and 8, and memory of 3, 2, 2, 1 and 1 for 331 Hz, 166 Hz, 100 Hz, 83 Hz and 66Hz, respectively).

### Diffusion analysis

For each trajectory that passed the filtering steps, the cumulative probability distribution of a particle’s displacement during a lag time of four frames (*t* = 12 ms) was calculated (except in Fig. 2a, where *t* = 3 ms was used to resolve multiple mobility components, if present). This lag time was chosen to reduce the influence of motion blurring, which is often referred to as a dynamic measurement error^25^, on our measurements of particle displacement. This dynamic error results in an underestimation of particle displacements at short lag times, which we observed at lag times below 12 ms (Supplementary Fig. 13a). To calculate diffusion coefficients we fitted the following one-, two-, and three-component models to the calculated cumulative probability distribution, *P*(*r, t*)^5^:

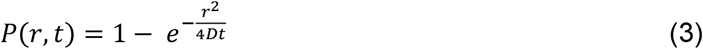

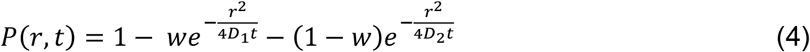

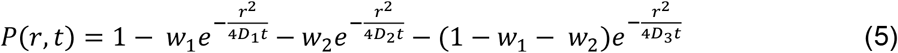

where *r* represents particle displacement during the chosen lag time, *t; D1, D2*, and *D3* represent the mobility components and *w1* and *w2* their weightings (summing up to 1). A trajectory was characterised as having more than one mobility component if adding an additional component improved the mean squared residual of the fit by more than one order of magnitude. Using this criterion, < 5% of trajectories measured in this study displayed two mobility components and none displayed 3 mobility components at t = 3 ms (Supplementary Fig. 7a). At *t* = 12 ms < 1% of trajectories displayed more than one mobility component. To characterise the diffusive behaviour of different oligomeric species, only trajectories that had a mean mass within two standard deviations of the mean mass of an oligomer were used (see Supplementary Fig. 8 and 11 for examples of this selection window). Additionally, trajectories that fit the following criteria were excluded from diffusion analysis:

- More than 20% of the trajectory points were gaps
- The mean of all contrast values of a trajectory differed by more than 20% from the value determined by Gaussian fitting
- The trajectory was too stationary (characterised by having a fast and slow mobility component and a weighting factor of less than 0.3 on the fast component).

For example, in the WT dataset shown in Fig. 2a, among 315 trajectories used in the analysis, 6 trajectories had too many gaps, 2 were too stationary and 2 had mean contrasts that differed significantly form the trajectory contrast determined by Gaussian fitting. These criteria helped eliminate trajectories that were strongly influenced by background fluctuations or a result of incorrect trajectory linking (Supplementary Fig. 22). Using this approach, histograms of the diffusion coefficients were plotted for each oligomeric species (Supplementary Fig. 7b, 9, 12) and the mean diffusion coefficient of each oligomer was calculated by fitting a Gaussian to these distributions. For the small number of trajectories that displayed two diffusion components, only the major component was included in these histograms. The number of histogram bins was determined using the Freedman-Diaconis rule.

### Residence time analysis

To calculate the dissociation rate constants from the SLB of each oligomeric species, we slightly modified the trajectory filtering procedure described above to limit the impact of background noise on measured trajectory lengths. Firstly, we constructed spatial maps to identify regions that were either particularly noisy (high standard deviations in pixel intensity in raw frames) or showed anomalously high particle densities (often a sign of improperly bound particles or increased background features). Trajectories that contained at least ten points (>33 ms in length) outside of these identified regions were kept and filtered using the standard deviation of their contrast as determined by Gaussian fitting, in a similar way as described above. Trajectories that contained less than 20 contrast measurements could sometimes result in poor Gaussian fits and in this case the median of the trajectory’s contrast measurements was used as the trajectory contrast. Additionally, trajectories that had coordinates within 5 pixels of the edge of the field of view were discarded. After these filtering steps, contrast histograms (70 bins) were plotted and trajectories were sorted by oligomeric species as described above. The distribution of trajectory lengths of a given oligomeric species was fit to the probability density function of a 1-component exponential distribution and the rate parameter was optimised by maximum likelihood estimation. To correct for the threshold of 33 ms (10 frames) that was applied in prior filtering steps, it was necessary to scale the probability density function by incorporating the threshold as an additional parameter:

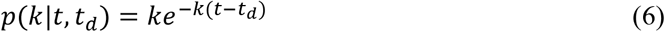

where *k* is the dissociation rate constant from the membrane, *t*_*d*_ is the time threshold applied during filtering, and *t* the trajectory length. This process was repeated for each oligomeric species detected in the dynamic MP measurements of WT (7 repeats) and ΔPRD (4 repeats) taken at 10-20 nM to achieve satisfactory particle density on the SLB (Supplementary Fig. 14-15).

### GTP data analysis

For the analysis of the effect of GTP on the mass distribution of WT on the SLB (Fig. 2g) the trajectories were filtered as described in the trajectory analysis section. Mass histograms were plotted for the movie recorded immediately before GTP was added (last 1 min of acquisition before GTP addition) and the first movie recorded immediately after GTP addition (first minute of acquisition after addition). Trajectories were allocated to a particular oligomeric species if their mass, as determined by Gaussian fitting was within two standard deviations of the mean mass of that oligomer, which was determined from the entire movie series up to GTP addition in each case (Supplementary Fig. 8b-g). This procedure was used for five replicate measurements of WT-dynamin (10-20 nM to achieve satisfactory particle density) with GTP addition (1 mM) after 3-4 min of acquisition (Supplementary Fig. 17a-b). We used the same approach for the measurement in which the non-hydrolysable GTP analogue, GMPPNP, was added instead of GTP (Supplementary Fig. 17c-d).

## Data and code availability

All raw movies that were used in this manuscript, the corresponding background-subtracted dynamic MP movies and the data used in the figures are available on request. The python software used for image processing, and particle identification, fitting and trajectory linking is available on request along with jupyter notebooks that outline the use of this package, from raw movies to linked particle trajectories. We can also provide the jupyter notebooks that were used to extract mass distributions, diffusion coefficients and dissociation rate constants from the spreadsheet of linked trajectories, which includes all filtering steps as described in the methods section.

## Acknowledgements

We thank Samuel Tusk for helpful discussion, contributions to the analysis software and troubleshooting during development. Additionally, we thank Stephen Thorpe and Nikolas Hundt for further contributions to the software and Max Felix Hantke for development of the PSF model and fitting procedure. We also thank Samuel Tusk, Stephen Thorpe, Jack Peters and Lee Priest for maintenance of the mass photometry setups used in this work. P.K., is supported by an ERC Consolidator grant (PHOTOMASS 819593). E.D.B.F. was supported by the Engineering and Physical Sciences Research Council (EPSRC) and Medical Research Council (MRC) (EP/L016052/1), St Hugh’s College and the Clarendon Fund. We thank Catherine Lichten for feedback on the manuscript.

## Author contributions

Concept: M.S.K., G.Y., P.K. Methodology: all authors Software: E.D.B.F., G.Y. Investigation: M.S.K., E.D.B.F., G.Y. Formal analysis: E.D.B.F., G.Y. Original draft: E.D.B.F., M.S.K., P.K. Review and editing: all authors Visualisation: E.D.B.F., P.K., M.S.K. Supervision: P.K.

