## supplementary information for "Label-free tracking and mass measurement of single proteins on lipid bilayers"

Eric D. B. Foley[#†], Manish S. Kushwah[#†], Gavin Young[†‡] and Philipp Kukura[†*]

[#]Equal contribution

[†]Physical and Theoretical Chemistry Laboratory, Department of Chemistry, University of Oxford, UK

[‡]Refeyn Ltd., Oxford, Oxfordshire, GB

### Contents

Supplementary Figure 1 ... 1

Background subtraction in standard MP vs dynamic MP

Supplementary Figure 2 ... 2

Effect of background subtraction on baseline noise

Supplementary Figure 3 ... 3

Effect of filtering particle trajectories by their SLB residence times

Supplementary Figure 4 ... 4

Contrast vs mass calibration of dynamic MP and standard MP

Supplementary Figure 5 ... 5

Mass distribution of WT on the SLB vs in solution

Supplementary Figure 6 ... 6

Effect of frame averaging on particle contrast

Supplementary Figure 7 ... 7

Extraction of diffusion coefficients from trajectories

Supplementary Figure 8 ... 8

Mass distributions of WT by dynamic MP

Supplementary Figure 9 ... 9

Distributions of diffusion coefficients of WT oligomers

Supplementary Figure 10 ... 10

Relationship between the diffusion coefficient and the number of subunits of WT oligomers

Supplementary Figure 11 ... 11

Mass distribution of ∆PRD measurements by dynamic MP

Supplementary Figure 12 ... 12

Distribution of diffusion coefficients of ∆PRD oligomers

Supplementary Figure 13 ... 13

Effect of chosen lag time on calculation of diffusion coefficients

Supplementary Figure 14 ... 14

Distribution of trajectory residence times of ∆PRD oligomers on the SLB

Supplementary Figure 15 ... 15

Distribution of trajectory residence times of WT oligomers on the SLB

Supplementary Figure 16 ... 16

Relationship between the dissociation constant from the SLB and number of subunits of WT oligomers

Supplementary Figure 17 ... 17

Effect of GTP addition on the mass distribution of WT

Supplementary Figure 18 ... 18

Custom-built setup used in this study

Supplementary Figure 19 ... 19

Effect of window size of median background subtraction on particle contrast

Supplementary Figure 20 ... 20

Image processing for identification of particle candidates
Supplementary Movie 1
Supplementary Movie 2
Supplementary Movie 3
Supplementary Movie 4
References ................................................................................................................ 23

#### Supplementary Figure 1

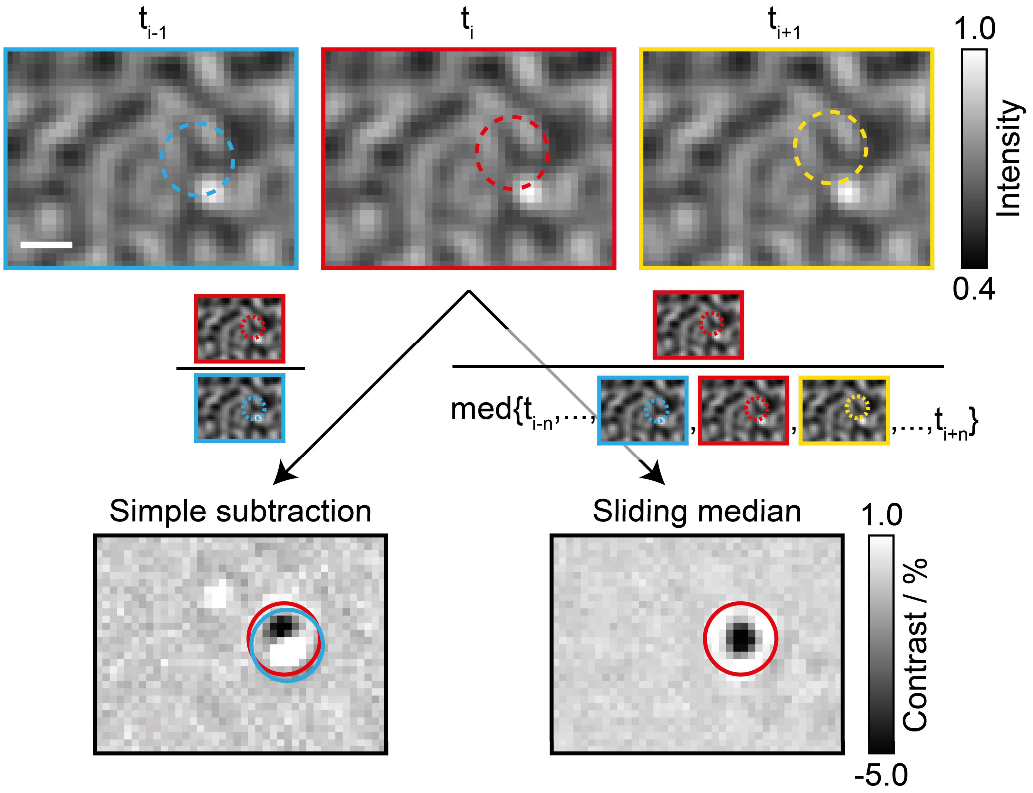

**Background subtraction in standard MP vs dynamic MP**

Zoom on three consecutive raw images containing a particle diffusing upwards through the image, which is masked by the large signal from the glass surface. In standard MP (left), images are divided by preceding images to remove the large static signal due to surface roughness[1]. This background subtraction relies on small particles binding to the surface and afterwards remaining stationary to visualise them. When particles bind to and diffuse on the surface, the background subtraction used in standard MP results in a signal that is a convolution of the particle's position at $t_{i-1}$ and $t_i$, which is challenging to reliably detect and quantify. A sliding median filter (right), *i.e.* subtracting each image's temporal median background obtained from a defined window of images around the image of interest, reveals signals of only the particles in the image of interest ($t_i$). For further explanation see 'Data processing' in the methods section. Scale bar: 500 nm.

#### Supplementary Figure 2

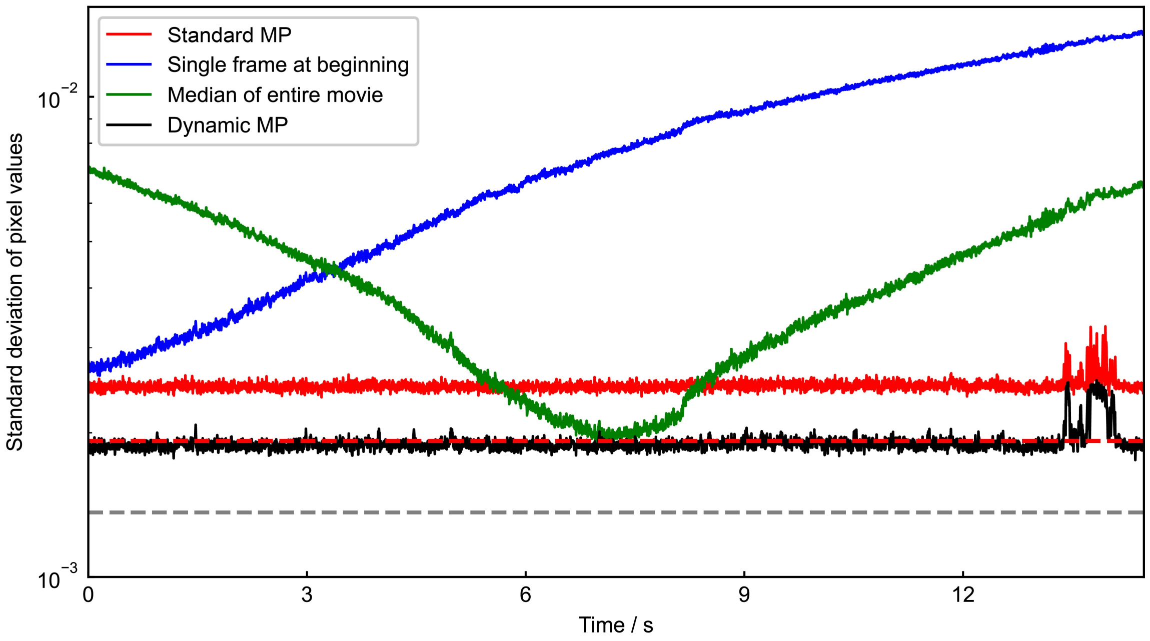

**Effect of background subtraction on baseline noise**

The baseline noise (the standard deviation of all pixel values in a given frame) of a dynamic MP movie of buffer solution on an SLB obtained with the background subtraction used in standard MP (red line), by subtracting the initial frame from all frames (blue line), subtracting the median of the entire movie from all frames (green line) and using the sliding median filter used for the dynamic MP experiments in this work (black line). The grey dashed line shows the theoretical shot noise limit when using the sliding median filter. The shot noise limit when using standard MP background subtraction (red dashed line) overlaps with the sliding median baseline noise (~0.002). The sliding median background subtraction results in lower background noise because the background is calculated from 200 additional frames compared to the background subtraction used in standard MP. Occasional spikes in background noise appear due to larger particles diffusing across the field of view.

#### Supplementary Figure 3

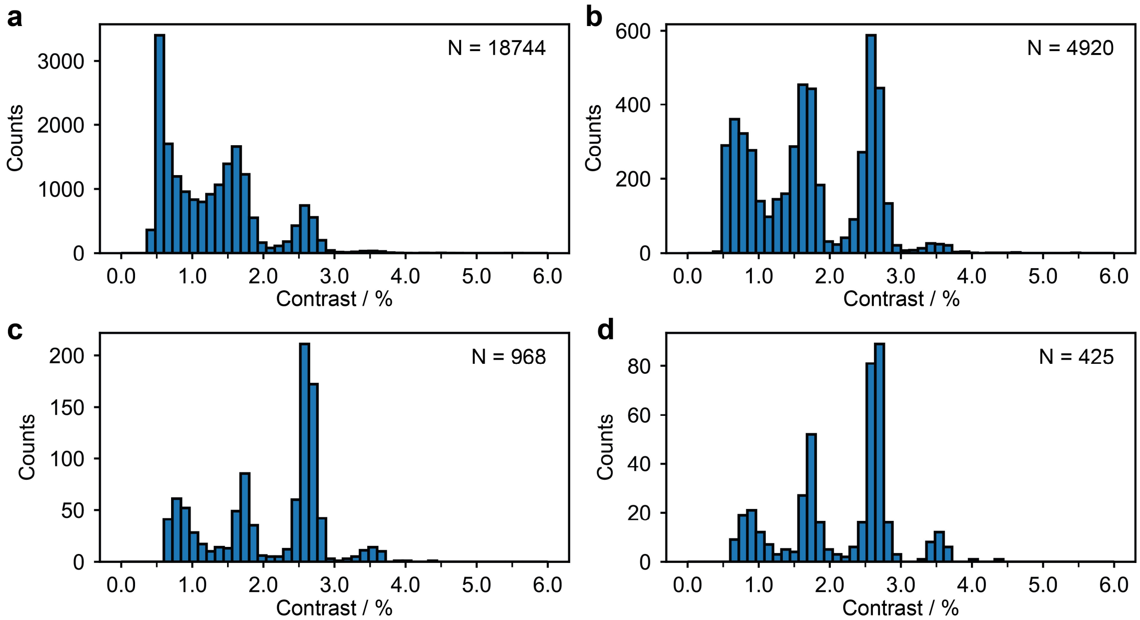

**Effect of filtering particle trajectories by their SLB residence times**

**(a)-(d)** Contrast histograms obtained from the same dynamic MP movie of WT used in Fig. 1b-d and 2a after applying a minimum threshold of 5, 10, 30 and 50 frames (331 Hz) to the trajectory length, respectively. Particle trajectories that remained on the membrane for fewer frames than the specified threshold were discarded. Here, each data point corresponds to the median contrast of a trajectory. This length filtering procedure effectively improves the quality of the data and reduces background features that were incorrectly identified as particles and linked into short trajectories, resulting in an increase in contrast resolution.

#### Supplementary Figure 4

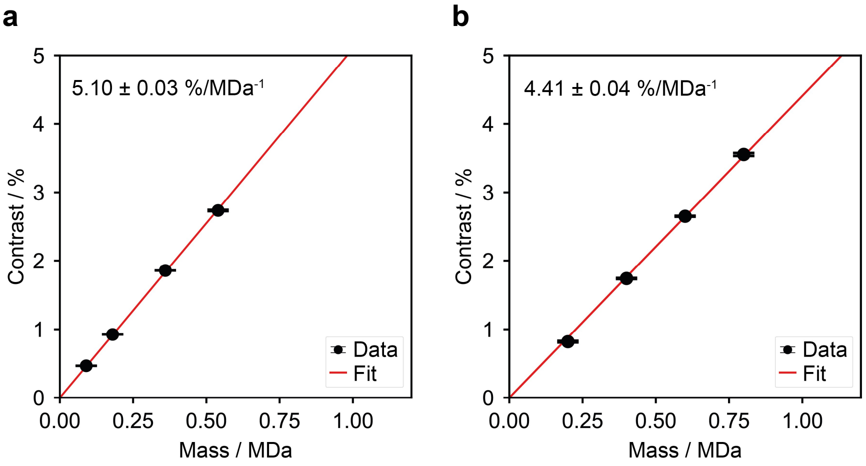

**Contrast vs mass calibration of dynamic MP and standard MP**

Contrast vs mass calibrations obtained from one measurement of ΔPRD (20 nM) in standard MP (**a**) and from dynamic MP measurements on an SLB in contact with WT (20 nM) taken within the next hour (Fig. 1b) (**b**). The contrast to mass calibration measured in dynamic MP movies was consistently 10-15% lower than that measured in standard MP. This effect is likely a result of particle motion during image acquisition, which results in motion blurring of the PSF. This effect increased as we increased frame averaging (Supplementary Fig. 6). Error bars represent the standard error of the mean of the Gaussian contrast distribution of each oligomeric species. The standard errors in contrast were generally < 0.02%, which causes the caps of the error bars to overlap. We used ΔPRD in standard MP because it spans more oligomeric states than WT under our experimental conditions.

#### Supplementary Figure 5

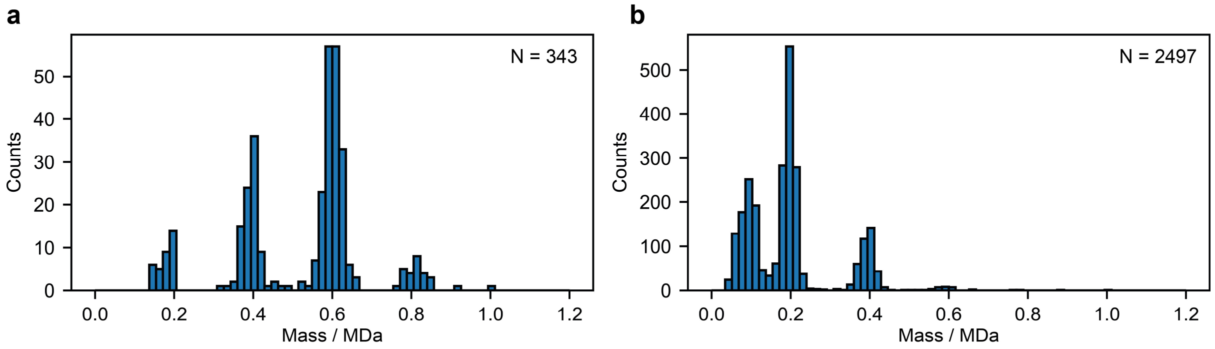

**Mass distribution of WT on the SLB vs in solution**

**(a)** Mass distribution of 20 nM WT in HKS-100 buffer (see methods) diffusing on an SLB. This data is the same as the one shown in Fig. 1b after additional filtering steps (see methods). **(b)** Three combined repeat measurements of 20 nM WT in HKS-100 buffer measured by standard MP. The peaks represent WT monomers (0.1 MDa), dimers (0.2 MDa), tetramers (0.4 MDa), hexamers (0.6 MDa) and octamers (0.8 MDa). The standard MP measurements were acquired at 191 Hz and then processed at a final integration time ~31 ms (effective frame rate ~32 Hz). The dynamic MP measurement was acquired at 331 Hz to minimise motion blurring (see main text), which resulted in higher baseline noise compared to the standard MP measurement and consequently WT monomers could not be detected.

#### Supplementary Figure 6

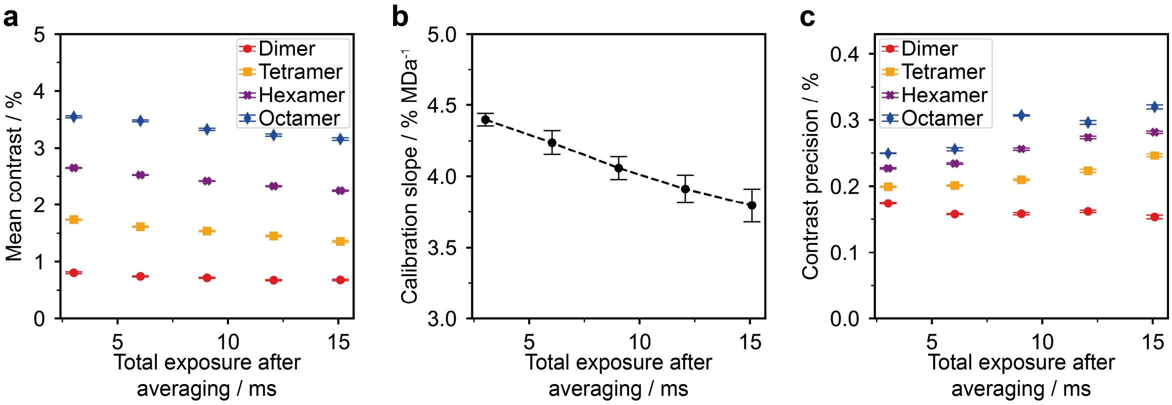

**Effect of frame averaging on particle contrast**

**(a)** Mean contrast of WT dimer (red circles), tetramer (orange squares), hexamer (purple crosses) and octamer (blue diamonds) trajectories vs single frame length after averaging. **(b)** Calibration slope obtained from the dynamic MP movie in (a) vs single frame length after averaging. **(c)** Contrast precision of our PSF-fitting procedure vs single frame length after averaging for each oligomer (same symbols as used in (a)). These effects are most likely a result of particle motion during image acquisition, which becomes more pronounced as more raw images are averaged together and the frame length increases. The plots were obtained from the same movie of WT used in Fig. 1b-d and 2a with additional frame averaging of 1, 2, 3, 4 and 5 frames, which corresponds to frame lengths of 3.02, 6.04, 9.05, 12.07 and 15.09 ms or frame rates of 331, 166, 110, 83 and 66 Hz, respectively. The error bars in (a) and (c) represent the standard error of the Gaussian fit to the distributions of the corresponding values of each oligomeric species. The error bars in (b) represent the standard deviation of the calibration slope. These plots were produced from the same set of dynamic MP movies used in Fig 1b-d and 2a (4 sets of 1 min movies of the same sample of 20 nM WT).

#### Supplementary Figure 7

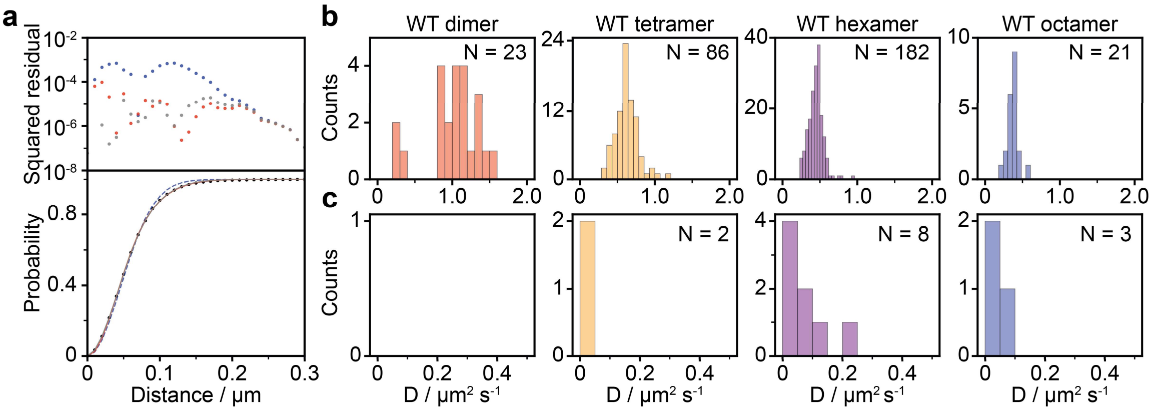

**Extraction of diffusion coefficients from trajectories**

**(a)** 1-(blue), 2-(red) and 3-(gray) component fits to the cumulative probability density of the distance moved during one frame (3 ms) by the particle shown in Fig. 1e-g in blue, red and grey, respectively. **(b)** Resulting distributions of the major diffusion component of WT dimer (red), tetramer (orange), hexamer (purple) and octamer (blue) particles for the same data used in Fig. 1 b-d and 2a (4 min movie of the same sample). **(c)** Same as **(b)** for the minor diffusion component (if present). The vast majority of particles only exhibited one diffusion component.

#### Supplementary Figure 8

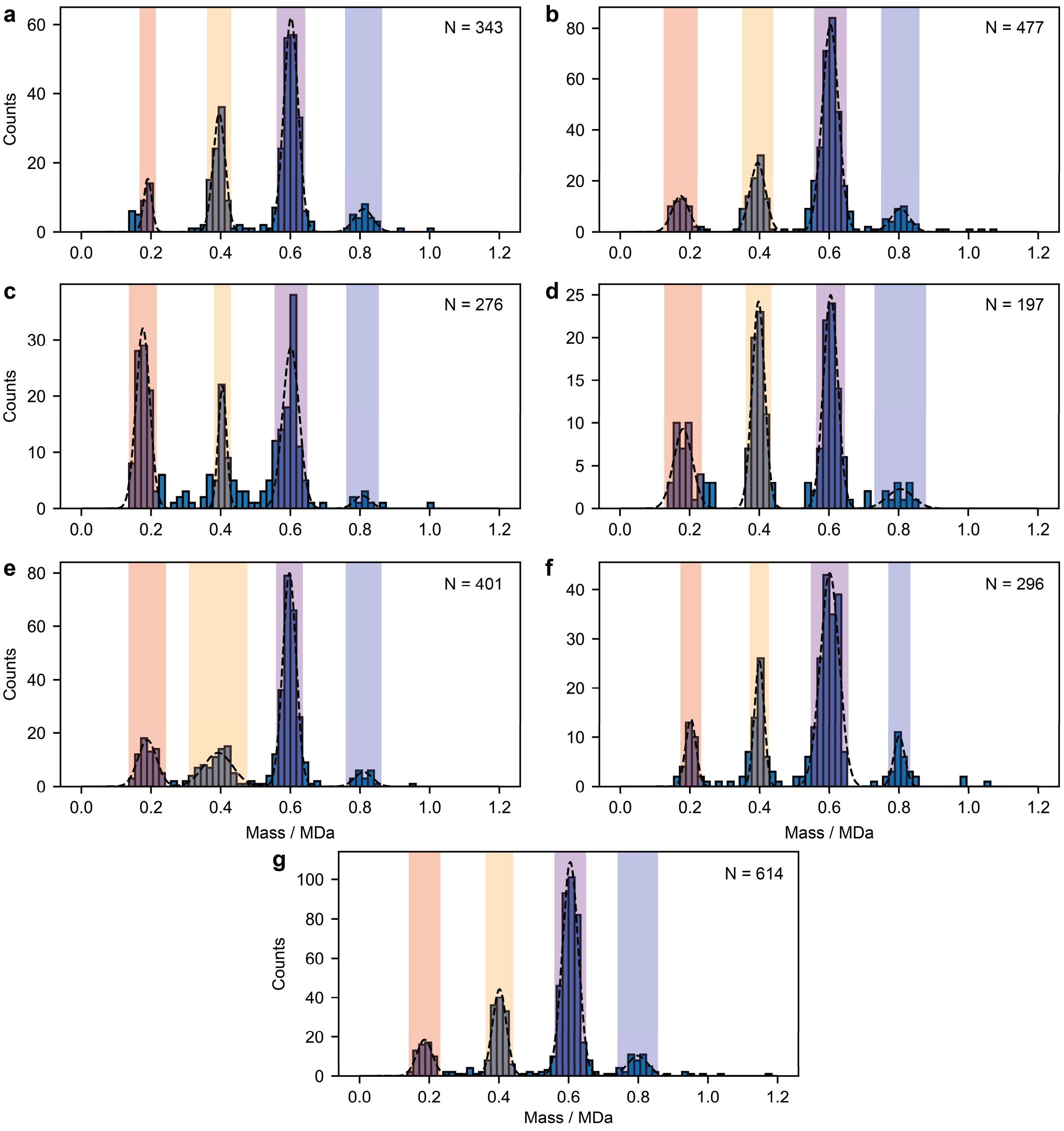

**Mass distributions of WT by dynamic MP**

**(a)-(g)** Mass histograms of seven repeats of dynamic MP measurements (4-5 min each) of WT (10-20 nM) in contact with an SLB. Each data point in the histograms represents the mean contrast of a trajectory determined by Gaussian fitting, as described in the methods section. The black dashed lines represent Gaussian fits to the oligomeric species of WT. If a trajectory's mean contrast fell within two standard deviations of the mean of these Gaussian fits, it was classified as that particular oligomeric species (red = dimer, orange = tetramer, purple = hexamer, blue = octamer). The number of bins for all mass histograms was set to 70 and the same initial guess for Gaussian fitting was used in each case. The dimer trajectories often overlapped in mass with background features (see also Supplementary Fig. 11) and were thus difficult to distinguish from background noise. As such, we excluded the WT dimer trajectories in the analysis of diffusion coefficients and residence times.

#### Supplementary Figure 9

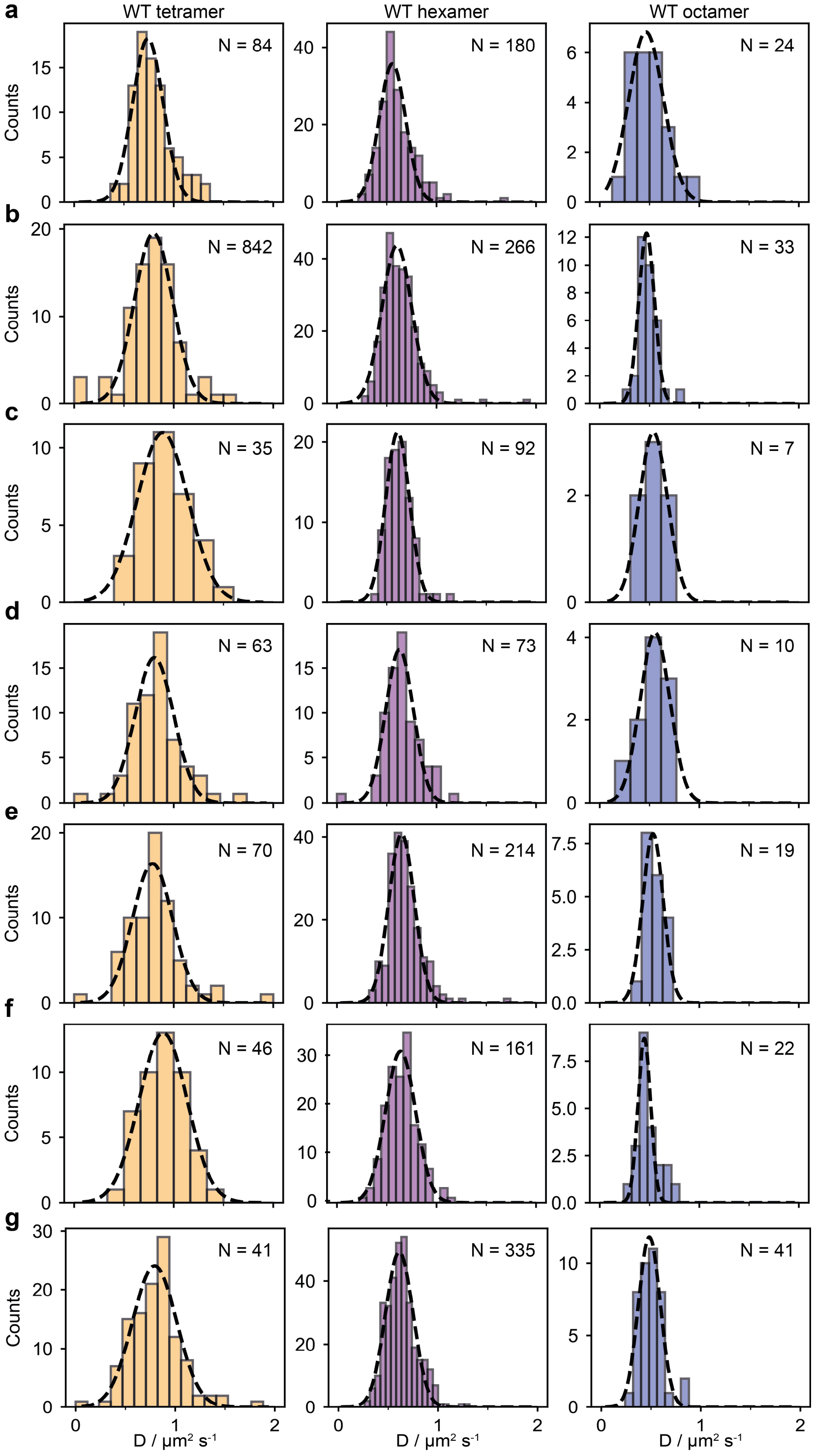

**Distributions of diffusion coefficients of WT oligomers**

**(a)-(g)** Diffusion coefficients calculated as described in the methods section for each oligomeric species (tetramer = orange, hexamer = purple, octamer = blue) detected in the WT measurements shown in Supplementary Fig. 8. The mean diffusion coefficient was determined by Gaussian fitting to the histograms (black dashed line). The number of bins in each histogram was chosen the Freedman-Diaconis rule.

#### Supplementary Figure 10

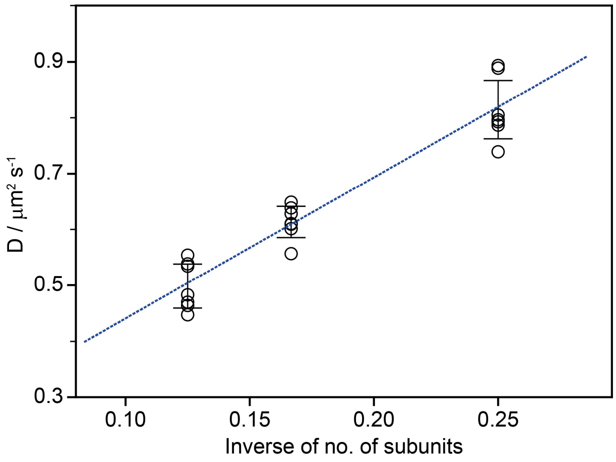

**Relationship between the diffusion coefficient and the number of subunits of WT oligomers**

Diffusion coefficients of each oligomeric species of WT from data shown in Supplementary Fig. 9 vs the inverse of the number of subunits of each oligomeric species and a corresponding weighted linear fit (blue dashed line). The error bars represent the standard deviation in the diffusion coefficients of each oligomeric species.

#### Supplementary Figure 11

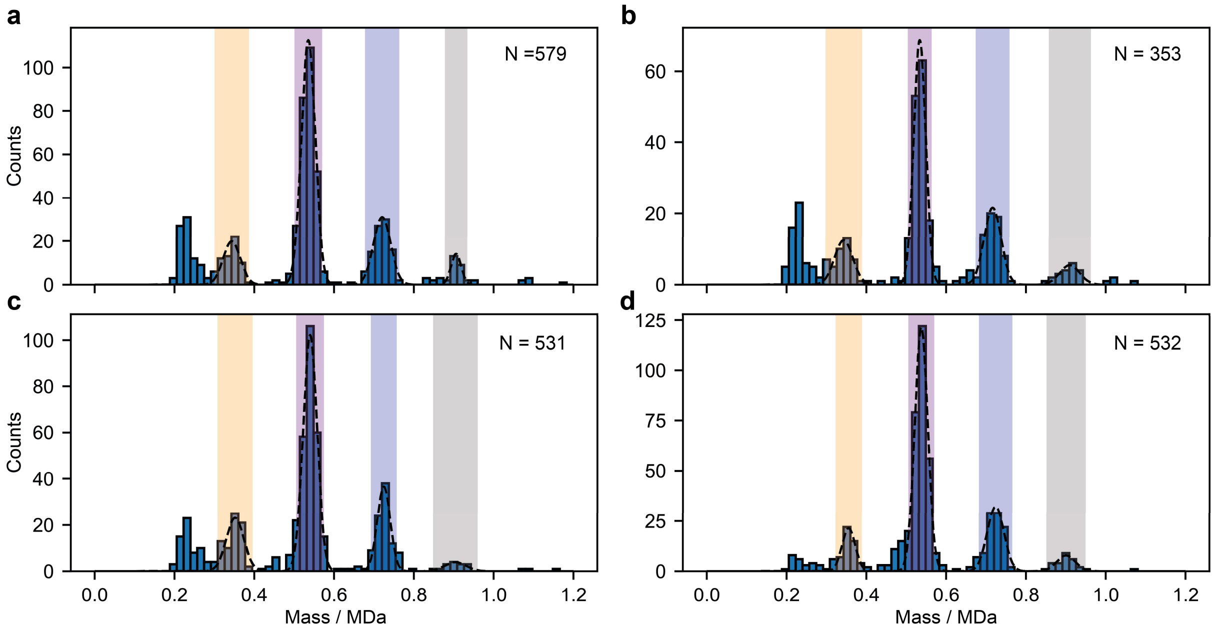

**Mass distribution of ΔPRD measurements by dynamic MP**

**(a)-(d)** Four repeats of dynamic MP measurements (4-5 min each) of ΔPRD in contact with an SLB. Each data point in the histograms represents the mean contrast of a trajectory determined by Gaussian fitting, as described in the methods section. The black dashed lines represent Gaussian fits to the oligomeric peaks of ΔPRD. If a trajectory's mean contrast fell within two standard deviations of the mean of these Gaussian fits, it was classified as that particular oligomeric species (orange = tetramer, purple = hexamer, blue = octamer, grey = decamer). The number of bins in all histograms was set to 75 and the same initial guess for Gaussian fitting was used in each case. The peak just above 0.2 MDa is a result of background noise and represents the limit of detection in these measurements. As such, the ΔPRD dimer (180 kDa) could not be reliably detected.

#### Supplementary Figure 12

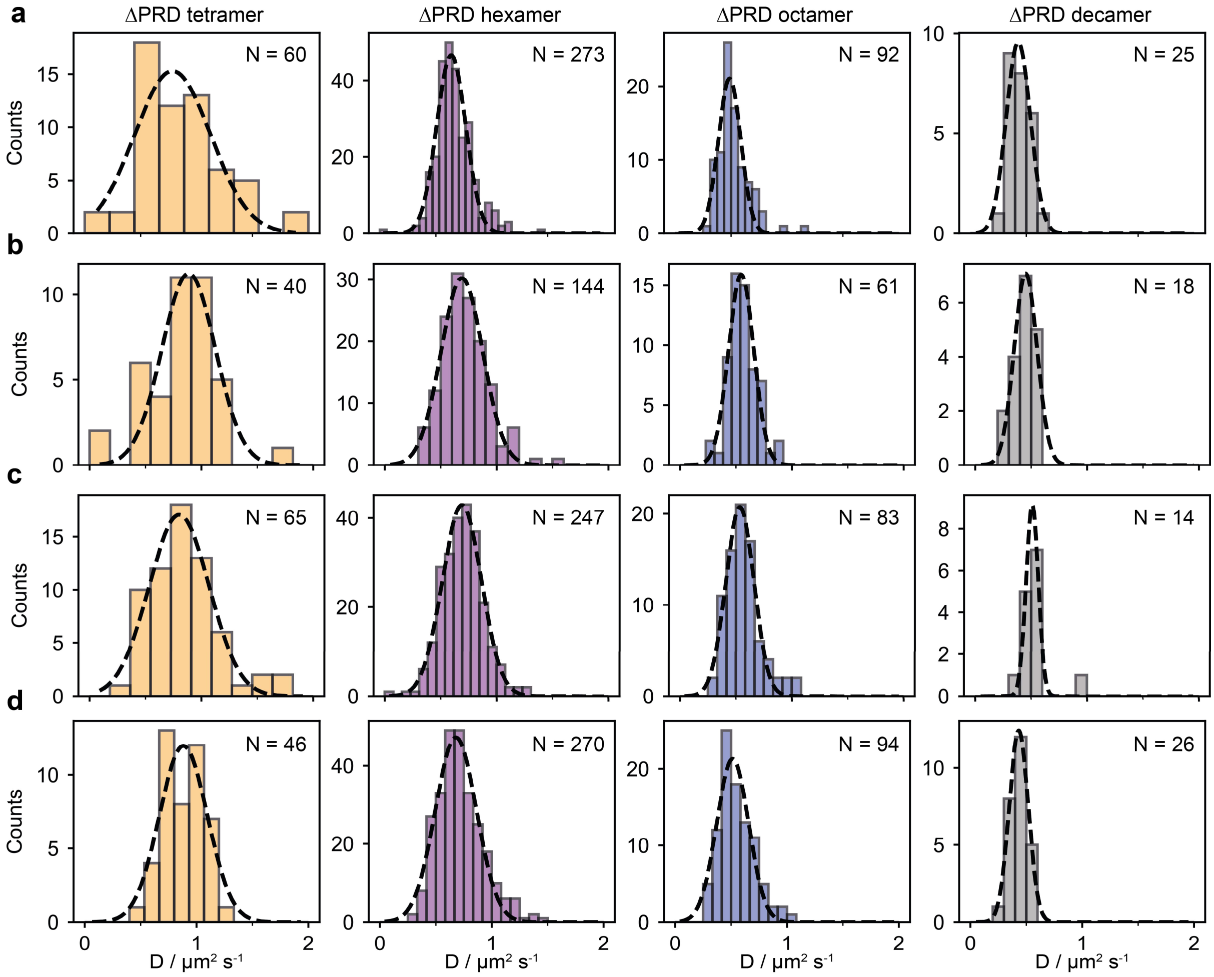

**Distribution of diffusion coefficients of ΔPRD oligomers**

**(a)-(d)** Diffusion coefficients calculated as described in the methods section for each oligomeric species (tetramer = orange, hexamer = purple, octamer = blue, decamer = gray) of the ΔPRD measurements shown in Supplementary Fig. 11. The mean diffusion coefficient was determined by Gaussian fitting (black dashed line) to the histograms. The number of bins in each histogram was chosen using the Freedman-Diaconis rule.

#### Supplementary Figure 13

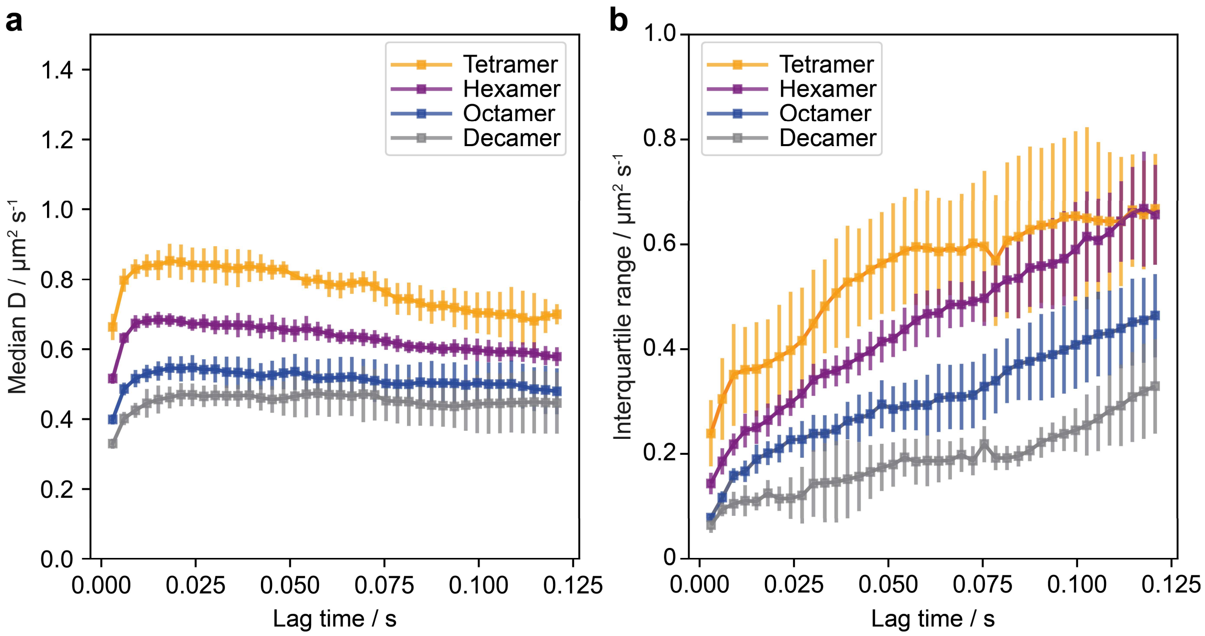

**Effect of chosen lag time on calculation of diffusion coefficients**

(**a**) Average median diffusion coefficient and **(b)** corresponding interquartile range vs chosen lag time (*t*) for each oligomeric species of the ΔPRD measurements shown in Supplementary Fig. 11-12. As the distribution of diffusion coefficients broadened significantly as the lag time increased, the diffusion coefficient of each oligomer was determined by taking the median of the distribution. Each data point represents the mean diffusion coefficient from the median values determined from four repeats of ΔPRD measurements and the error bars show the standard deviation.

#### Supplementary Figure 14

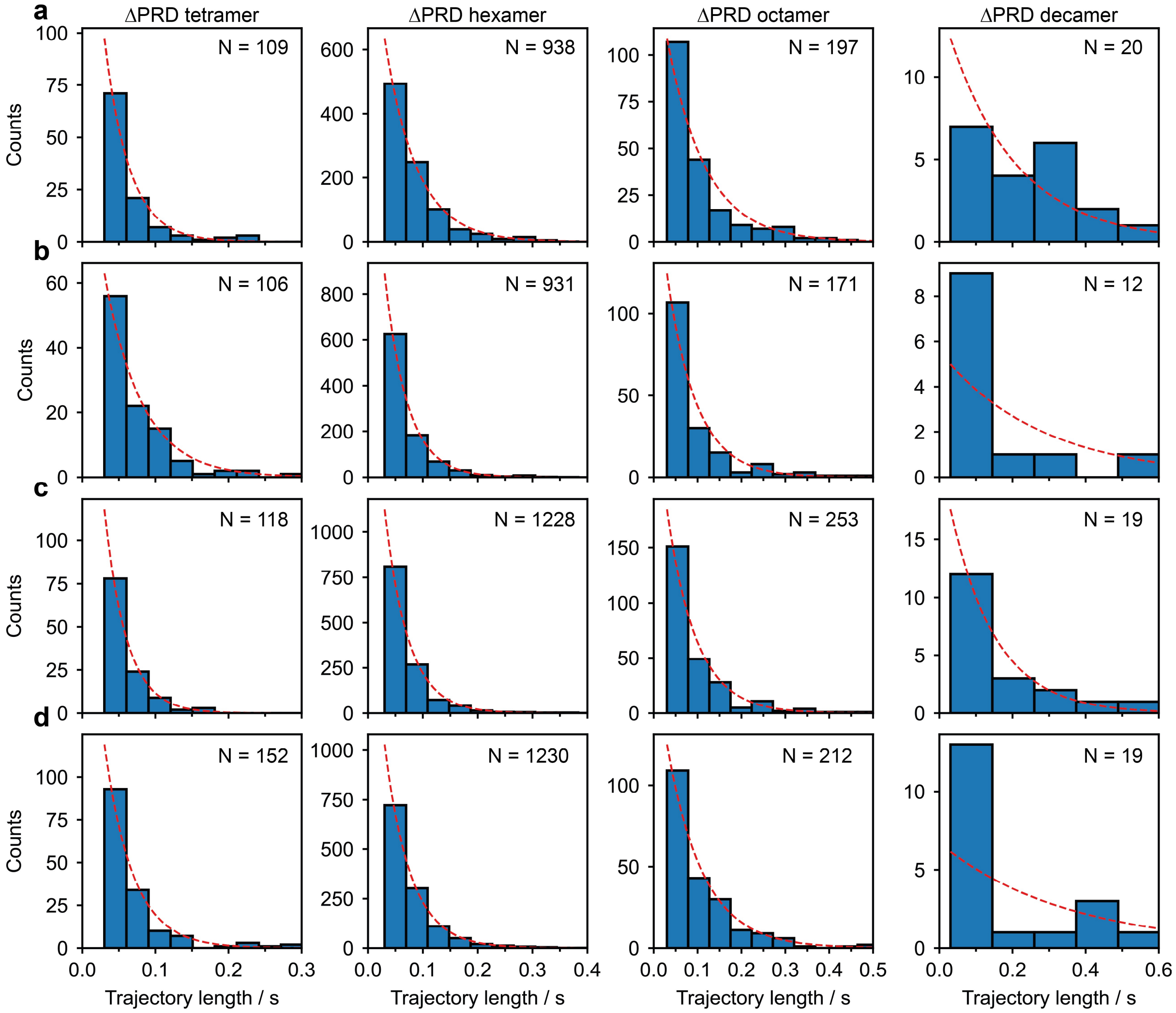

**Distribution of trajectory residence times of ΔPRD oligomers on the SLB**

**(a)-(d)** Histograms of counts vs trajectory lengths for each oligomeric species detected in the four repeats of ΔPRD measurements shown in Supplementary Fig. 11-12. Dissociation rate constants for unbinding from the SLB were calculated by maximum likelihood estimation using the unbinned data (see methods) and the resulting exponential distributions are plotted as red-dashed lines (appropriately scaled for display). The number of decamer trajectories (right column) was small in comparison to the other oligomeric species and thus sometimes resulted in poor fits.

#### Supplementary Figure 15

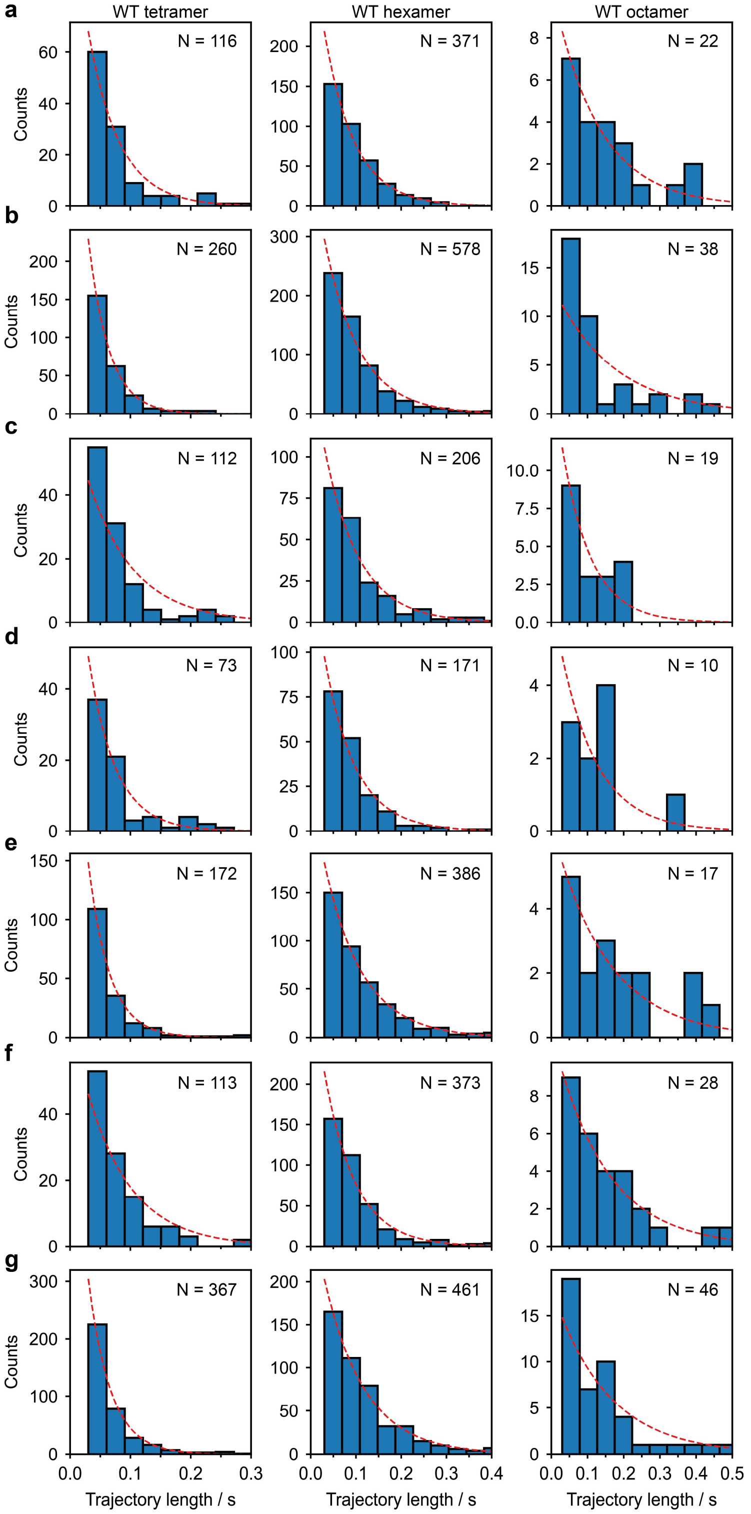

**Distribution of trajectory residence times of WT oligomers on the SLB**

**(a)-(d)** Counts vs trajectory length for each oligomeric species detected in the 7 replicate measurements of WT shown in Supplementary Fig. 8-9. Dissociation rate constants for unbinding from the SLB were calculated by maximum likelihood estimation using the unbinned data (see methods) and the resulting exponential distributions are plotted as red-dashed lines (appropriately scaled for display). The number of octamer trajectories (right column) was small in comparison to the other oligomeric species and thus sometimes resulted in poor fits.

#### Supplementary Figure 16

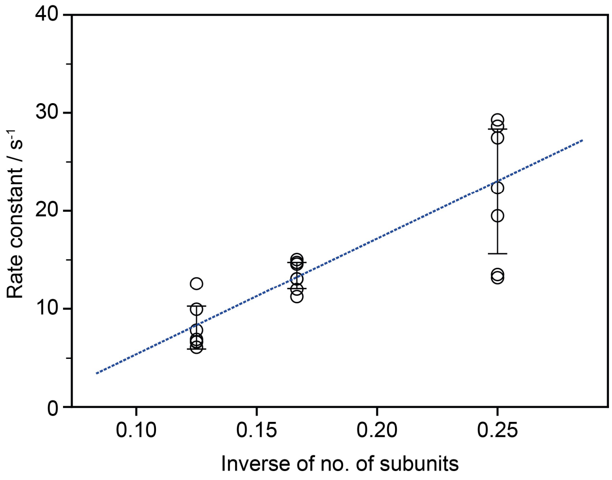

**Relationship between the dissociation constant from the SLB and number of subunits of WT oligomers**

Dissociation constant from the SLB of each oligomeric species of WT determined from the data shown in Supplementary Fig. 15 vs the inverse of the number of subunits of each oligomeric species and a corresponding weighted linear fit (blue dashed line). The error bars represent the standard deviation in the dissociation constants of each oligomeric species determined from seven repeat measurements.

#### Supplementary Figure 17

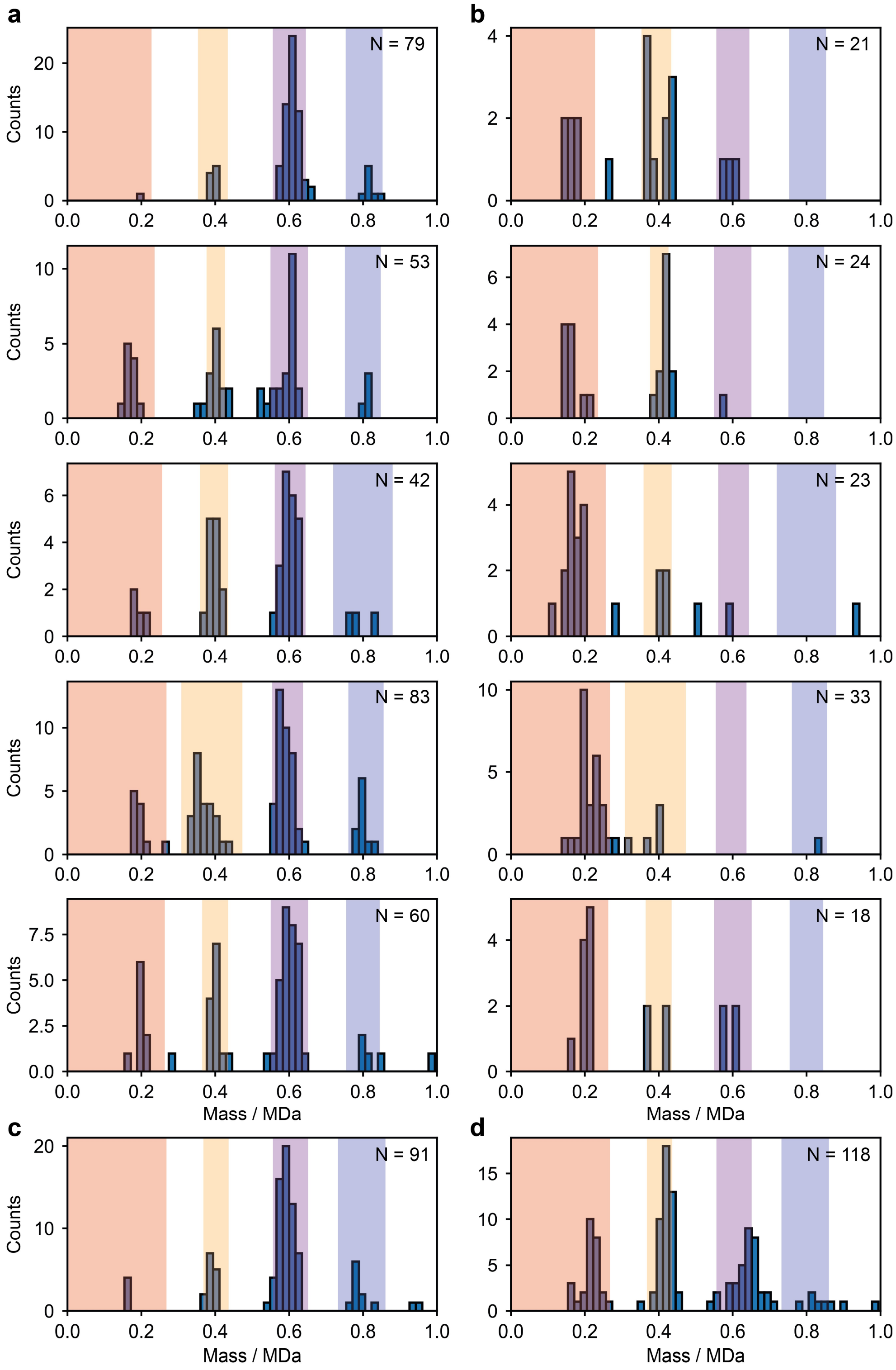

**Effect of GTP addition on the mass distribution of WT**

Mass distribution of five measurements of WT (10-20 nM) in contact with an SLB before (**a**) and after (**b**) addition of 1 mM GTP (1 min movie each). Measurement of WT immediately before **(c)** and after **(d)** addition of the non-hydrolysable GTP-analogue, GMPPNP. Shaded areas were used to classify trajectories as dimer (red), tetramer (orange), hexamer (purple) and octamer (blue) as described in the methods section. This data was used to generate the bar plot shown in Fig. 2g.

#### Supplementary Figure 18

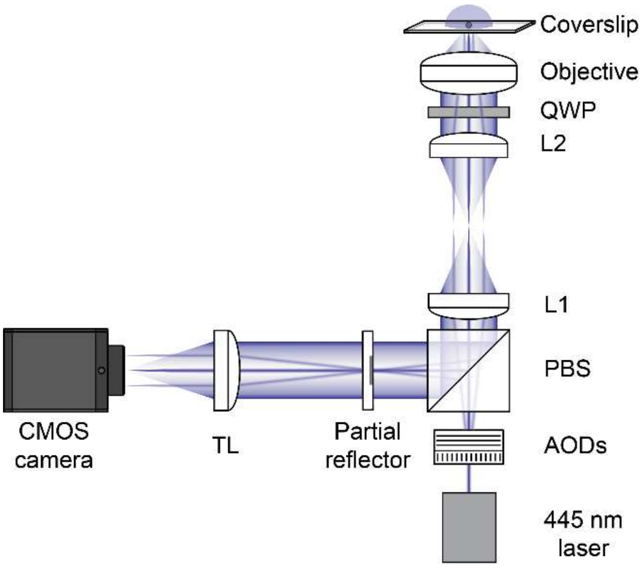

**Custom-built setup used in this study**

Layout of the custom-built mass photometer used to acquire the data shown in Fig. 1a, e-g and 2e-f, similar to that reported previously[1,2]. A collimated 445 nm laser beam (Lasertack) is directed through an orthogonal pair of acousto-optic deflectors (AODs; AA opto Electronic, DT SXY-400). After passing through a polarising beam splitter (PBS), two telecentric lenses (L1 and L2) direct the deflected beam through a quarter-wave-plate (QWP) and into the back focal plane of the objective (Olympus PlanApo N, 1.42 NA, 60x). This setup results in the beam being weakly focused and scanned across the sample to illuminate a 9.4 x 6.2 $\mu m^2$ field of view. The light reflected at the glass-water interface of the coverslip as well as the light scattered by the sample is collected by the objective and travels through the same telecentric lens system that the incident light was passed through. Use of the QWP and PBS, separates the collected light from the incident light and passes through a partial reflector, which is positioned in the reimaged back focal plane of the objective. This partial reflector is made of a 3.5 mm diameter layer of silver deposited onto a window and attenuates the reflective (at low NA) light by over two orders of magnitude compared to the light scattered by particles[1]. After passing through the partial reflector the light is imaged by a tube lens (TL) onto a CMOS camera (Point Grey, GS3-U3-23S6M-C) resulting in 250x magnification and a pixel size of 23.4 nm. Images are recorded at 1 kHz and then binned in groups of 3 by a custom LabVIEW software that also performs x and y pixel binning (generally 3 x 3) before saving the images for later analysis. Samples are mounted on a custom-built sample stage, with height adjustment provided by a micrometer translation stage (Optosigma) and piezoelectric actuator (Thorlabs AE0505D16F). Sample focus is maintained by an autofocus system (not shown) as follows: a diode laser (Lasertack) is coupled out of a single mode fiber with a 4x objective, and travels collimated into the imaging objective via a dichroic. The beam overfills the objective back aperture and high-NA components are totally internally reflected, causing the back-reflection of an annular beam from the sample. This annular beam is imaged onto a CMOS camera (Thorlabs DCC1545M ). Custom labview code continuously measures the radius of this annular beam, and adjusts the piezoelectric actuator in the stage to maintain this radius at a set value. The sample stage and all the optics (except for coupling of the autofocus laser into the single mode fiber) are coupled to to a 600x400x50mm aluminium breadboard, enclosed by 40mm thick aluminium walls and lid, which sits on a granite table.

#### Supplementary Figure 19

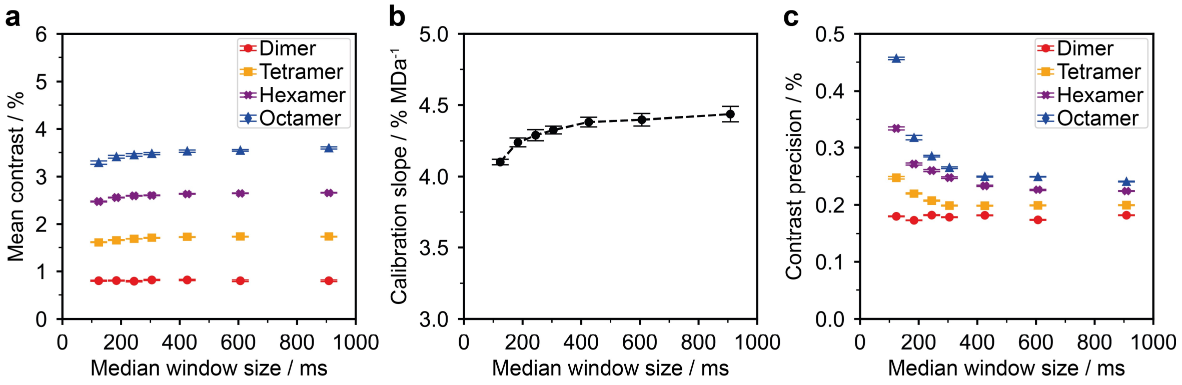

**Effect of window size of median background subtraction on particle contrast**

**(a)** Mean contrast of WT dimer (red circles), tetramer (orange squares), hexamer (purple crosses), octamer (blue triangles) trajectories vs window size chosen for the sliding median background subtraction (at 331 Hz). **(b)** Calibration slope obtained from the dynamic MP movie in (a) vs median window size. **(c)** Contrast precision of our PSF-fitting procedure vs median window size for each oligomer (same symbols as used in (a)). At small window sizes, the PSF of slow-moving, large particles may become part of the subtracted background if they move an insufficient distance during the time window included in the median filter (<2-3 pixels). The plots were obtained from the same movie of WT used in Fig. 1b-d and 2 a. The error bars in (a) and (c) represent the standard error of the Gaussian distributions of the corresponding values of each oligomeric species. The error bars in (b) represent the standard deviation of the calibration slope.

#### Supplementary Figure 20

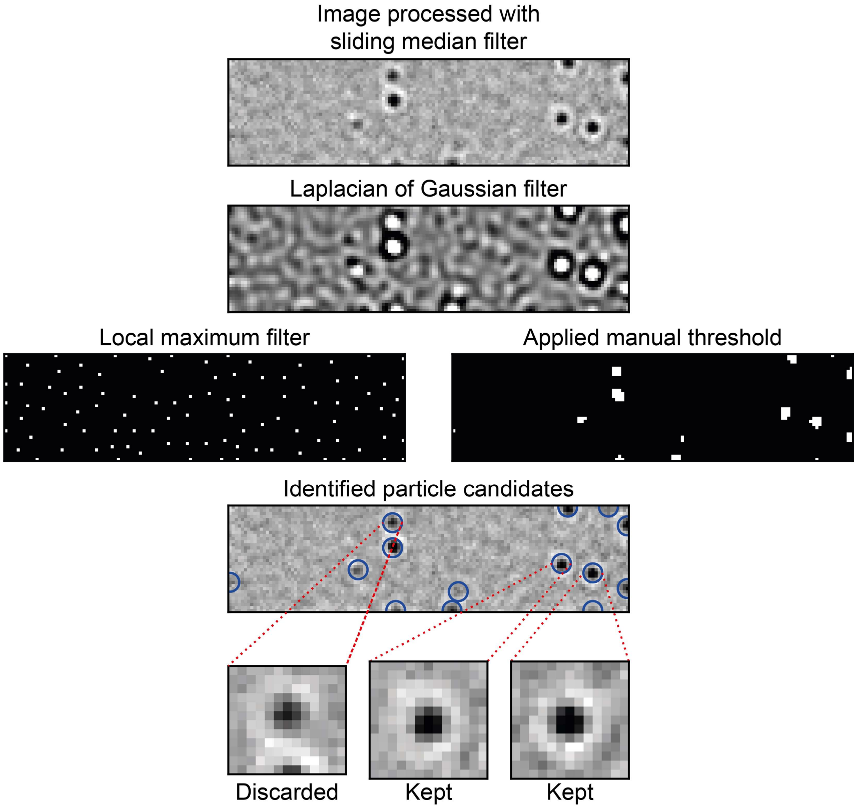

**Image processing for identification of particle candidates**

Background-subtracted images are filtered with a Laplacian of Gaussian kernel with a size to match the size of PSFs in the images (sigma = 1.5). Next, a manually set threshold and local maximum filter are applied to the filtered image. By combining the resulting binary maps, we obtain single pixels at the centre of potential particle candidates (circled in blue). From each of these pixels a 13x13 region of interest is constructed to which our PSF model is the applied for quantification of particle contrast and position. If the pixel candidate is too close to an edge of the field of view to construct a complete 13x13 region of interest, it is discarded, as shown in the example here.

#### Supplementary Figure 21

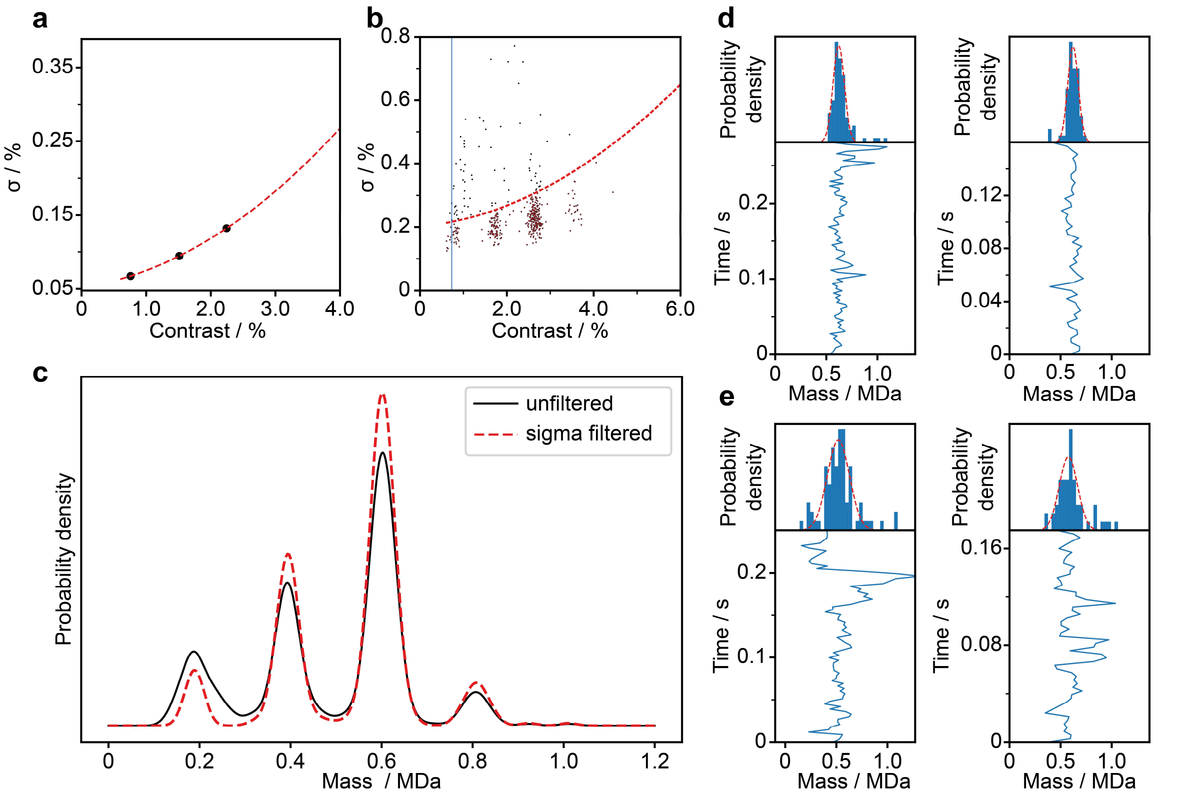

**Filtering of trajectories using the standard deviation of their contrast distributions**

**(a)** Standard deviation (σ) vs mass for WT oligomers obtained from standard MP experiments in solution. **(b)** Standard deviation vs mass scatter plot for the trajectories obtained from the dynamic MP measurement of WT shown in Fig. 1b-d and 2a (after length filtering). The solution-phase trend from (a) is applied with an appropriate offset as a threshold. Trajectories below the threshold (red) are used for analysis. In this particular case, trajectories with a standard deviation in mass below 170 kDa (blue line) were not used to reduce the number of trajectories that likely result from background noise in the analysis. **(c)** Effect of the applied threshold on the mass distribution of trajectories. **(d)** Examples of trajectories below the threshold (top panel, kept) and above the threshold (bottom panel, rejected).

#### Supplementary Figure 22

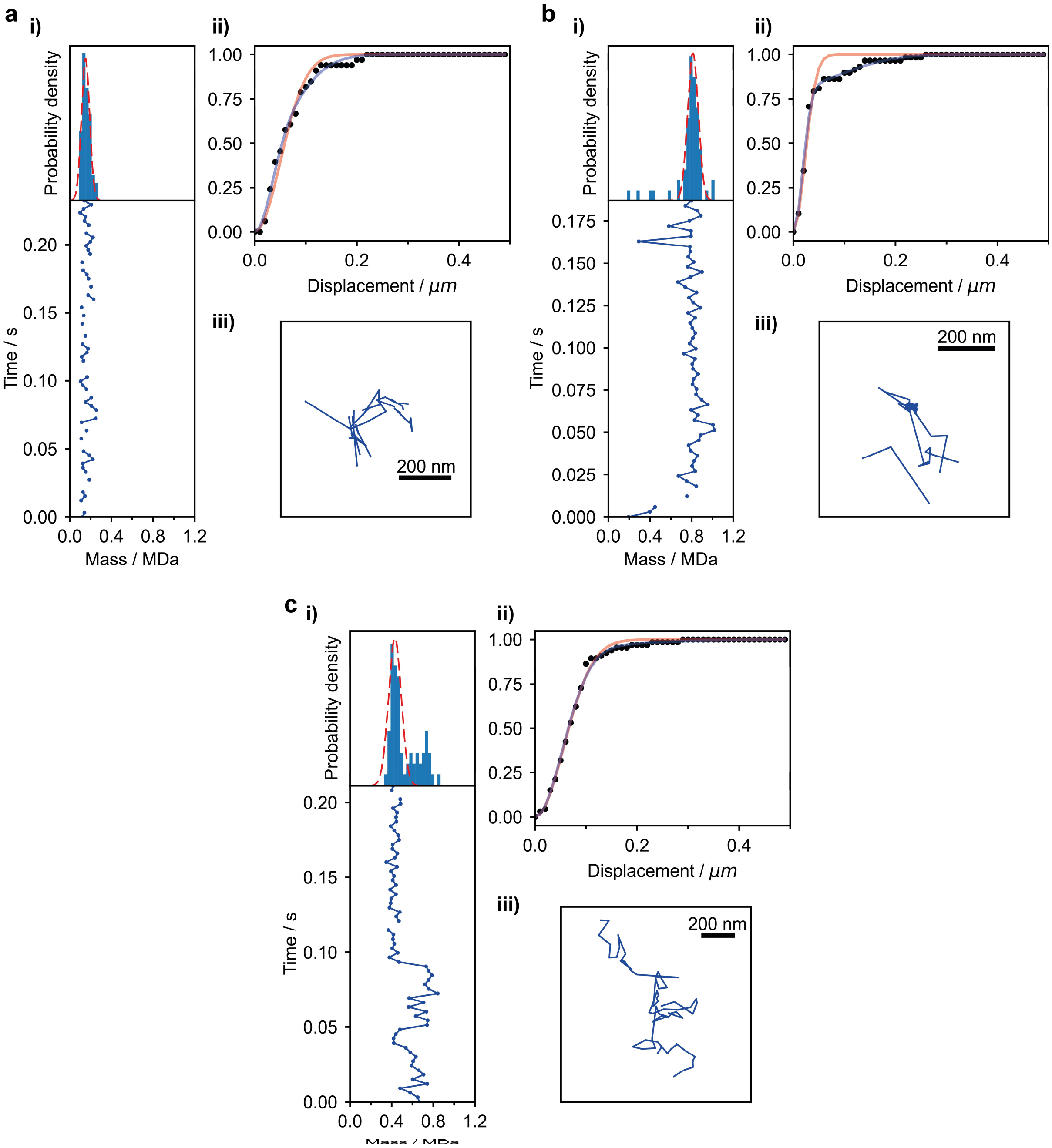

**Examples of particles that were excluded from the diffusion analysis**

**(a)** Mass trace (**i**), cumulative probability density distribution of 1-frame displacement with 1-and 2-component mobility fits in red and blue lines, respectively (**ii**), and trajectory in 2D space (**iii**) of a particle that was excluded from the diffusion analysis because it was above the set threshold of allowed trajectory gaps. (**b**) Same as in (a) but for a particle that was excluded because it was determined to be too stationary during its time on the SLB. **(c)** same as (a) but for a particle whose mean mass deviated too far from the mass determined by the Gaussian fitting algorithm (red dashed line), which was usually a sign of an incorrectly linked trajectory. These examples of excluded particles were chosen from the same data shown in Fig. 1b-c and 2a.

#### Supplementary Movies

##### Supplementary Movie 1

15 s section of the dynamic MP movie (displayed at 25 frames per second) of WT (20 nM) in contact with an SLB, corresponding to the data shown in Fig. 1b-c and 2a. Red dots represent locations of signals that were successfully identified as particles, quantified and linked into trajectories without applying a minimum residence time threshold. Scale bar equals 1 μm.

##### Supplementary Movie 2

Section (frames 3797-9974) of a dynamic MP movie of WT in contact with an SLB, in which the particle trajectory shown in Fig. 1e-g was identified (circled in red). This particle frequently approaches the edge of the field of view, which intermittently prevented detection and quantification by our processing software (indicated by the red circle briefly disappearing), and led to its trajectory being split up into several shorter ones. As a result, the short sub-trajectories had to be combined into the particle's entire trajectory by inspection. In this experiment, 6 μl of 1 μM WT was added to 60 μl of buffer on the SLB but with minimal mixing, which led to a gradual increase in particle density over time. Scale bar equals 1 μm. The movie is displayed at 25 frames per second.

##### Supplementary Movie 3

Zoom-in on a later section (frames 15900-16300) of the dynamic MP movie shown in Supplementary Movie 2, showing the dissociation event displayed in Fig. 2e. Scale bar equals 1 μm. The movie is displayed at 25 frames per second.

##### Supplementary Movie 4

Zoom-in on a section (frames 22500-22800) of a dynamic MP movie of ΔPRD (10 nM) in contact with an SLB, showing the association event displayed in Fig. 2f. Scale bar equals 1 μm. The movie is displayed at 25 frames per second.
